# An integrative approach to protein sequence design through multiobjective optimization

**DOI:** 10.1101/2024.03.01.582670

**Authors:** Lu Hong, Tanja Kortemme

## Abstract

With recent methodological advances in the field of computational protein design, in particular those based on deep learning, there is an increasing need for frameworks that allow for coherent, direct integration of different models and objective functions into the generative design process. Here we demonstrate how evolutionary multiobjective optimization techniques can be adapted to provide such an approach. With the established Non-dominated Sorting Genetic Algorithm II (NSGA-II) as the optimization framework, we use AlphaFold2 and ProteinMPNN confidence metrics to define the objective space, and a mutation operator composed of ESM-1v and ProteinMPNN to rank and then redesign the least favorable positions. Using the multistate design problem of the foldswitching protein RfaH as an in-depth case study, we show that the evolutionary multiobjective optimization approach leads to significant reduction in the bias and variance in RfaH native sequence recovery, compared to a direct application of ProteinMPNN. We suggest that this improvement is due to three factors: (i) the use of an informative mutation operator that accelerates the sequence space exploration, (ii) the parallel, iterative design process inherent to the genetic algorithm that improves upon the ProteinMPNN autoregressive sequence decoding scheme, and (iii) the explicit approximation of the Pareto front that leads to optimal design candidates representing diverse tradeoff conditions. We anticipate this approach to be readily adaptable to different models and broadly relevant for protein design tasks with complex specifications.

**Author summary:** Proteins are the fundamental building blocks of life, and engineering them has broad applications in medicine and biotechnology. Computational methods that seek to model and predict the protein sequence-structure-function relationship have seen significant advancement from the recent development in deep learning. As more models become available, it remains an open question how to effectively combine them into a coherent computational design approach. One approach is to perform computational design with one model, and filter the design candidates with the others. In this work, we demonstrate how an optimization algorithm inspired by evolution can be adapted to provide an alternative framework that outperforms the post hoc filtering approach, especially for problems with multiple competing design specifications. Such a framework may enable researchers to more effectively integrate the strengths of different modeling approaches to tackle more complex design problems.

## Introduction

The field of computational protein design has achieved major breakthroughs in recent years [1] in terms of its ability to design proteins and protein assemblies with diverse folds and functions (see, e.g., [2–9]), which has already found application in the design of therapeutically relevant biomolecules such as vaccines [10] and antibodies [11,12]. Such breakthroughs are built upon improvements in atomistic modeling techniques, such as the Rosetta software suite [13], and recent advances in machine learning-based structure prediction models [14–18], sequence design (or inverse folding) models [19–25], protein language models [26–34], and denoising diffusion probabilistic models [35–44].

As more models become available and the space of designable protein machinery becomes more sophisticated, there is an increasing need for frameworks that can integrate multiple models and objective functions directly into the protein design process, in order to take advantage of the strengths of different modeling approaches and to ensure that the designs satisfy all the desired structural, biophysical, enzymatic, and/or therapeutical specifications. One way this can be achieved is through post hoc filtering, whereby a generative model is used to propose designs, which are then scored using one or more different models to screen for designs with desired characteristics. The success of this strategy critically depends on the degree of overlap between the region of the design space explored by the generative model, and the regions of the design space favored by the filters. A low degree of overlap leads to a high rejection rate and likely sub-optimal design candidates.

A common cause of low overlap is that the filters conflict with each other or the generative model, and thus cannot be simultaneously optimized. This type of scenario is prevalent in protein design; for example, foldswitching proteins (also known as metamorphic proteins [45]), or more generally, proteins that undergo conformational changes necessary for their function, may require, at a given position, different sidechain hydrophobicities in the different conformational states, resulting in the impossibility of simultaneous optimization of the folding free energies of all relevant states. To fully account for such tradeoff conditions requires an explicit approximation of the Pareto front in the objective space. A key property of the Pareto front is that the solutions on the Pareto front dominate those outside of the Pareto front; more precisely, for any feasible solution *B* outside of the Pareto front, one can find a solution *A* on the Pareto front that is no worse than *B* on all objectives and improves *B* on at least one objective. In other words, the Pareto front represents the optimal tradeoff conditions among the objective functions, and thus should be considered the optimization target when multiple models and objective functions are needed to specify the design problem. Although it is straightforward to perform a non-dominated sorting during post hoc filtering, there is in general no guarantee that the non-dominated solutions represent a close approximation or unbiased sampling of the Pareto-optimal solutions, especially when they are produced by a generative model not designed to optimize the objective functions.

In this work, we therefore seek an alternative, integrative framework for sequence design that can more closely guide the generative process and fully account for the Pareto front in the objective space. Specifically, we argue that evolutionary multiobjective optimization algorithms [46], a class of genetic algorithms designed for multiobjective optimization, provide one such suitable framework that can be adapted to meet these criteria. This is because these algorithms are, by construction, designed to explicitly approximate the Pareto front in a user-specified objective space, and, as we demonstrate in this work, can iteratively guide the sampling process through the construction of biophysically-informed mutation operators. This emphasis on both explicit approximation of the Pareto front and use of informative mutation operators distinguishes this work from previous applications of genetic algorithms to protein design [47–53].

Multistate design [54] provides a natural setting to demonstrate how an evolutionary multiobjective optimization framework might function in practice, because the need to explicitly represent multiple states of a protein in this type of design problems directly maps onto the ability of the optimization algorithm to simultaneously consider multiple objective functions. In particular, we perform a detailed analysis of the multistate sequence design problem for *E. coli* RfaH, using the established Non-dominated Sorting Genetic Algorithm II (NSGA-II) [55] as the optimization framework (Fig 1). RfaH is a difficult model system for multistate design because it is a foldswitching protein that undergoes extensive conformational changes between an N-terminal domain bound all-α (RfaHα) and a dissociated all-β (RfaHβ) C-terminal domain [56], which involves considerable differences in the secondary and tertiary structure and solvent accessibility of its amino acid residues. For the multistate design of RfaH, we have chosen three well-established models, each representative of a different modeling approach: the inverse folding model ProteinMPNN [19,20] (henceforth abbreviated as pMPNN), the protein language model ESM-1v [27], and the structure prediction model AlphaFold2 [14,57]. In particular, we use AlphaFold2-based confidence metrics to compute what is known as the AF2Rank composite score [58], and employ it as a measure of folding propensity, without the need for multiple sequence alignments.

**Fig 1.**
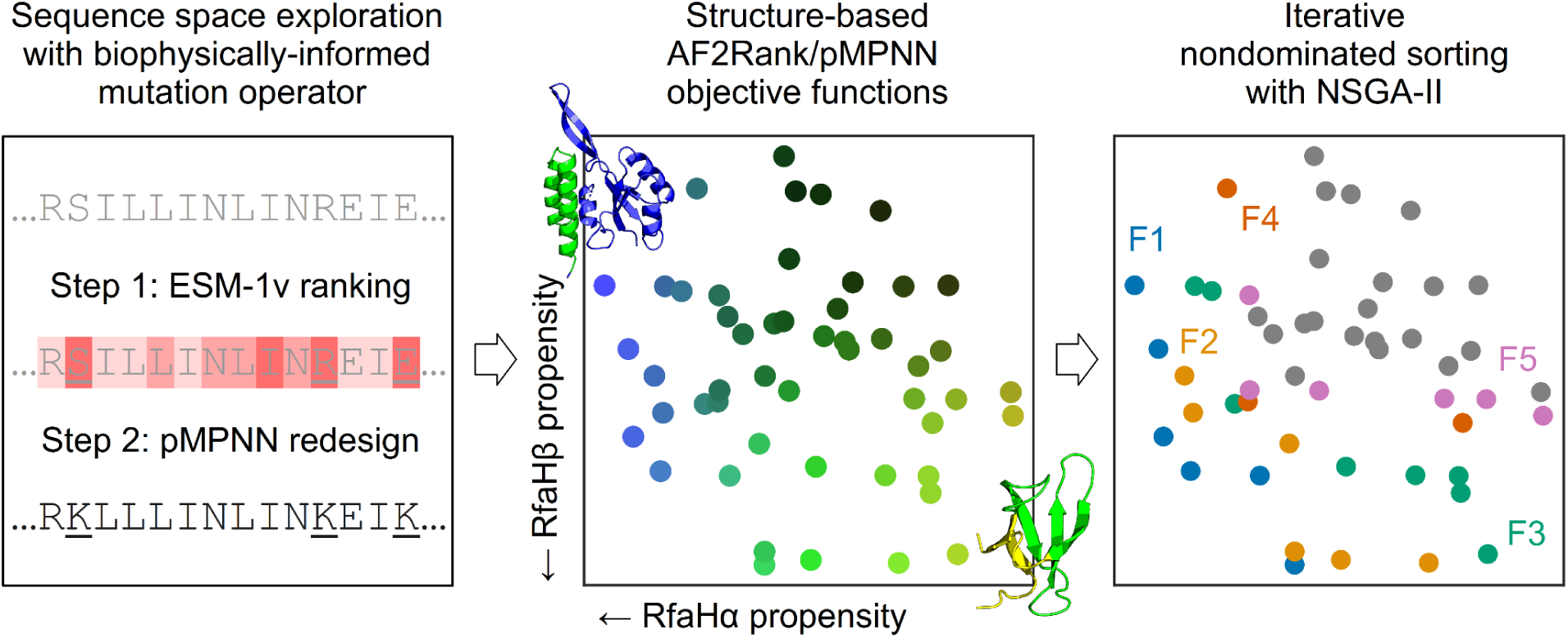
Evolutionary multiobjective optimization provides a suitable framework for multistate design. In this work, we examine how machine learning models such as pMPNN, AlphaFold2/AF2Rank, and ESM-1v may be integrated directly into protein sequence design through a multiobjective optimization method known as Non-dominated Sorting Genetic Algorithm II (NSGA-II). Left: first, new design candidates are proposed through a mutation operator; here, this operator is composed of ESM-1v, which is used to rank residue positions, and ProteinMPNN (pMPNN), which is used to redesign the least nativelike positions. Middle: the design candidates are then scored using objective functions derived from AlphaFold2 and pMPNN confidence metrics. Right: lastly, the scored candidates are sorted into successive pareto fronts (here numbered F1 to F5), and the candidates from the best fronts are selected by NSGA-II for the next round of design. To demonstrate the effectiveness of this framework, we perform an in-depth analysis of the multistate design problem for RfaH, a small foldswitching protein whose C-terminal domain can interconvert between an all-α RfaHα state and an all-β RfaHβ state. The designable positions (residue 119 to 154) are highlighted in green in the cartoon representations of the two RfaH states in the middle panel; note that the N-terminal domain is not shown in the ribbon representation of the RfaHβ state (see Methods).

Through our benchmark analysis, we find that NSGA-II, in conjunction with the basic random resetting mutation operator, is uncompetitive with a direct application of pMPNN. However, by embedding pMPNN and ESM-1v directly into the mutation operator, whereby ESM-1v is used to rank the designable positions and pMPNN is used to redesign the least nativelike residues, the genetic algorithm is able to better approximate the Pareto front in the objective spaces, which translates into designs with reduced sequence entropy and improvement in native sequence recovery, especially at positions where pMPNN alone fails. Together, these results indicate that evolutionary multiobjective optimization is a suitable framework for integrating multiple models directly into the protein sequence design process, and there is considerable value in constructing informative mutation operators to guide and accelerate the exploration of protein sequence space.

## Results

### The random resetting mutation operator results in slow convergence

In this work, we seek to assess whether multiobjective evolutionary optimization algorithms constitute a suitable framework for integrating multiple models and sources of information into protein sequence design, using the multistate design of the foldswitching protein RfaH as an in-depth test case. In general, such a genetic algorithm has three components (Fig 1): (i) a method for proposing new candidates, typically through the use of (binary) tournament selection, crossover operators, and mutation operators, (ii) a set of objective functions for scoring the proposed candidates, and (iii) an algorithm for forming a new population from the existing and proposed candidates. For the first two components, we will investigate how pMPNN, AF2Rank, and ESM-1v can be incorporated into sequence design through the mutation operators and objective functions (for the crossover operator, we find that the number of crossover points has a minimal impact on the design outcome (S2 Fig); therefore, for all following simulations, we set the number of crossover points to two and focus on the mutation operator instead). For the third component, we employ NSGA-II [55], an established genetic algorithm for multiobjective optimization (see Methods for more details).

In the following benchmark analysis, we will employ pMPNN both as an objective function and as a generative model in the context of both single-state and multistate design. For the sake of clarity, we will refer to pMPNN as pMPNN-SD (single-state design) when it is used to generate sequences conditioned on the structure of a single state, and pMPNN-AD (average decoding) when it is used to perform multistate design by averaging logits over multiple states during sequence decoding; when pMPNN is used as an objective function, we will refer to its log likelihood score for the full sequence, conditioned on the structure of a single state, as the pMPNN-SD log likelihood. In contrast, sequences designed using NSGA-II will always be referred to with the label GA (genetic algorithm), followed by a description of the simulation setup in square brackets.

We begin with the simplest baseline setup using the random resetting operator: starting with a population of fully randomized RfaH C-terminal domain sequences (residues 119 to 154), with a probability controlled by the mutation rate parameter, each designable position is selected for redesign, and the redesign is done with uniform sampling over the set of standard (20 naturally occurring) amino acids. The redesigned sequences are then scored against the RfaHα and RfaHβ state, with either the pMPNN-SD log likelihood or the AF2Rank composite score (see Methods). This setup is closely related to protein design via hallucination [59,60], and, in a sense, inverts the learned sequence-to-structure mapping of AlphaFold2; similar methods have been explored in the literature, typically with unsatisfactory results [20,21,43]. Consistent with the literature, we find that this baseline setup is slow to converge by all considered metrics (Fig 2, first two columns), fails to reach parity with pMPNN-AD (Fig 2, gray dotted lines), and the resulting sequence profiles exhibit a high degree of sequence entropy, especially for those designed using the AF2Rank objective functions (S3 Fig and S8 Fig), indicating that these simulations have not converged in the sequence space. These results suggest that a naive application of a genetic algorithm is likely to be outperformed by a single-pass inverse folding model, and AlphaFold2 is, on its own, a poor mutational effect predictor [58,61].

**Fig 2.**
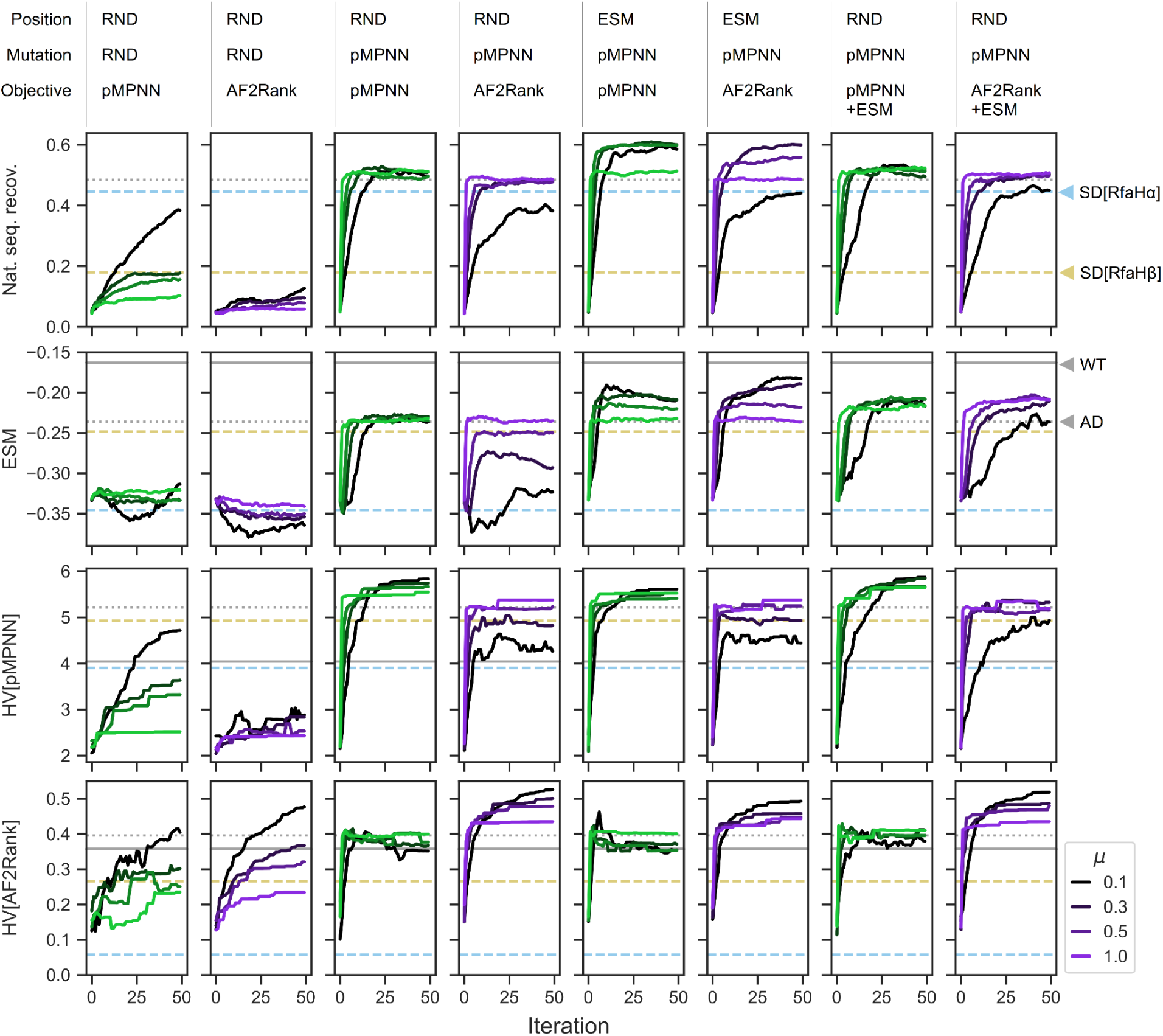
Integration of multiple models through evolutionary multiobjective optimization improves protein sequence design outcomes. Each column represents one genetic algorithm setup, indicated by the legend above each column that describes how, at each iteration, the subset of designable positions is selected, how the mtations are proposed, and which objective functions are used to score the designs; here, RND stands for random, pMPNN stands for ProteinMPNN, and ESM refers to ESM-1v. Each row represents a different quality metric: “nat. seq. recov.” refers to the population average fractional identity to the WT sequence, “ESM” refers to the ESM-1v log likelihood score, averaged over both the population and sequence positions, “HV” refers to hypervolume in the pMPNN-SD objective space (HV[pMPNN]), and hypervolume in the AF2Rank composite score objective space (HV[AF2Rank]); see Methods for more details. Each panel represents the progression of a genetic algorithm simulation over 50 iterations and at four different mutation rates (*μ*). Within each panel, the horizontal lines represent the quality metric values calculated for the WT RfaH sequence (solid gray), the population average of the pMPNN-AD design sequences (dotted gray), the population average of the RfaHα pMPNN-SD design sequences (dashed blue), and the population average of the RfaHβ pMPNN-SD design sequences (dashed yellow). Simulations results at additional mutation rates are shown in S1 Fig.

### A pMPNN-based mutation operator improves autoregressive sequence decoding

Next, we investigate whether the poor performance of the baseline setup can be rescued with a more informative mutation operator. From this point on, we will refer to each GA setup with three descriptors: how the subset of designable positions is selected, how the mutations are proposed, and how the designed candidates are scored; for example, the baseline setup with the random resetting operator and the AF2Rank objective functions can be denoted this way as GA[RND,RND,AF2Rank], where RND stands for random.

While the random resetting operator allows unbiased exploration of the sequence space, it is inefficient, given the combinatorial complexity of the protein sequence space. On the other hand, inverse folding models such as pMPNN have likely learned a lower-dimensional manifold in the sequence space, conditioned on the backbone conformation, that allows for efficient and accurate decoding of native-like sequences. Motivated by this observation, we modify the random resetting operator, so that the selected subset of designable positions are passed onto pMPNN-AD for redesign. We note here that this setup is similar to a recent method [53] that incorporated pMPNN into a genetic algorithm, although in [53], pMPNN is used in addition to, rather than in place of, the random resetting operator, and each application of pMPNN modifies all designable positions, rather than a subset thereof. With this modification, for at least some choices of mutation rate, the genetic algorithm is able to reach parity with pMPNN-AD in terms of native sequence recovery and the average ESM-1v log likelihood score, and outperforms pMPNN-AD in terms of hypervolume [62], a metric that assesses the optimality and diversity of candidate sequences (see Methods) (Fig 2, third and fourth columns; compare with the gray dotted lines).

The GA[RND,pMPNN,pMPNN] setup provides a useful case study in the potential advantages of genetic algorithms. This specific setup injects no additional biophysical information into the sequence design process, yet the resulting Pareto front shows clear separation in the pMPNN-SD objective space from the population generated with pMPNN-AD (S4 Fig). This result demonstrates the potential advantage of the evolutionary multiobjective optimization framework over the post hoc filtering approach to sequence design: although it is possible to approximate the Pareto front in the objective space by a post hoc non-dominated sorting from the pMPNN-AD population, the resulting approximate Pareto front will still be dominated by the design population generated by the genetic algorithm.

What could explain the performance difference between pMPNN-AD and GA[RND,pMPNN,pMPNN]? This difference is partly explained by the fact that a genetic algorithm is a numerical optimization technique, as compared to the autoregressive Monte Carlo sampling steps of pMPNN; as such, suboptimal residue types that are selected against in a genetic algorithm may be retained, even if only with low probability, by pMPNN-AD alone. More importantly, this difference rests on a basic limitation of autoregressive decoding: the first positions to be decoded tend to have the highest level of uncertainty, because more of the sequence context remains masked initially. Here, the genetic algorithm setup alleviates this problem in two ways: by reducing the number of masked positions in each pass, and by revisiting the same positions over the iterations, presumably with better sequence context each time. More precisely, with the GA[RND,pMPNN,pMPNN] setup, the number of redesigned positions with each application of the mutation operator and the number of times a position is redesigned both follow binomial distributions. This explanation is consistent with the observation that the gap between the GA[RND,pMPNN,pMPNN] sequences and the pMPNN-AD sequences narrows as the mutation rate increases (S4 Fig), because a higher mutation rate increases the number of masked positions in each pass, and it weakens the Markovian dependence between successive iterations.

### ESM-1v accelerates sequence space exploration

Lastly, we examine the effect of incorporating ESM-1v into the genetic algorithm. Given that ESM-1v is a good variant/mutational effect predictor [27], we propose two approaches to incorporating it into the design process. First, similar to a method described in [63] that prioritizes sequence design over low AlphaFold2 confidence regions, we use the ESM-1v marginal probabilities to rank and select the least nativelike positions for redesign (Fig 2, fifth and sixth columns); second, we directly incorporate the per-sequence average ESM-1v log likelihood score as a third objective function (Fig 2, last two columns). We find that the first approach (GA[ESM,pMPNN,pMPNN/AF2Rank]) produces the overall best performance, while the second approach (GA[RND,pMPNN,pMPNN/AF2Rank+ESM]) shows worse performance in terms of both native sequence recovery and average ESM-1v log likelihood score.

What could account for the performance differences between these two approaches? The difference in native sequence recovery can be rationalized by examining the pMPNN-SD design results (Fig 2): on average, the RfaHα single-state redesigned sequences are more similar to the pMPNN-AD sequences in terms of native sequence recovery, but the RfaHβ single-state redesigned sequences are more similar to the pMPNN-AD sequences in terms of the ESM-1v log likelihood score. As such, setting the ESM-1v score as an objective function biases sampling towards RfaHβ-like sequences with lower identity to the WT (S5 Fig). In contrast, in the GA[ESM,pMPNN,pMPNN/AF2Rank] setups, because ESM-1v is not used to generate mutations, the effect of its preference for the RfaHβ state is much weaker on native sequence recovery. In theory, the bias could have slowed down, or even stalled, convergence, by directing computation towards positions that are already nativelike, but not RfaHβ-like; however, comparison of the rate of convergence across the setups do not support this hypothesis (Fig 2).

The difference in the average ESM-1v log likelihood scores can be rationalized in terms of the structure of the simulation setups. In the second approach, optimization of ESM-1v as a third objective function is constrained by tradeoffs with the need to simultaneously optimize the pMPNN-SD or AF2Rank scores, which do not appear to be correlated with ESM-1v scores (S6 Fig). In contrast, in the first approach with the GA[ESM,pMPNN,pMPNN/AF2Rank] setups, because ESM-1v is used to set the design positions, the simulation prioritizes refinement of the lowest-ranked positions, which necessarily leads to an improvement in the sequence-averaged ESM-1v scores. As such, ESM-1v is invoked upstream of non-dominated sorting in the objective space, and the indirect optimization of its scores is not subject to the same tradeoff conditions in the second approach.

Interestingly, the effect of ESM-1v in the GA[ESM,pMPNN,AF2Rank] setup can be interpreted in the framework of Bayesian inference:

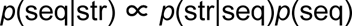

where the posterior distribution *p*(seq|str) is sampled by pMPNN-AD during sequence decoding, the likelihood function *p*(str|seq) is approximated by the AF2Rank composite score, and the prior *p*(seq) is partially set by the mutation operator. The random resetting operator is equivalent to a flat prior, and gives every designable position equal priority for redesign. The use of ESM-1v to rank and select positions for redesign can be interpreted as an informative prior that accelerates traversal in the sequence space.

### RfaH multistate design shows sequence tradeoff patterns based on solvent accessibility

A key feature of an evolutionary multiobjective optimization algorithm is that the solutions approximate the Pareto front in the objective space. To the extent that objective functions such as the pMPNN-SD log likelihood and the AF2Rank composite score correlate with desirable biophysical properties such as stability, this Pareto optimality condition necessarily needs to manifest in the sequence space as mutations that improve such properties for either or both states of RfaH. Therefore, in this and the following sections, we investigate in detail the differences in the sequence profiles generated by pMPNN-SD, pMPNN-AD, and the genetic algorithm, and the potential biophysical consequences of such differences. Among the genetic algorithm setups we have examined so far, GA[ESM,pMPNN,pMPNN/AF2Rank] give the best overall performance, especially at the intermediate mutation rate 0.3; as such, we will focus on the sequences generated in the last iteration of these two sets of simulations in this analysis. For the sake of brevity, we will simply refer to these two setups and their last iteration sequences as GA[pMPNN] and GA[AF2Rank].

To understand the distribution of designed sequences in the sequence space, we first seek a low dimensional representation of the RfaH C-terminal domain sequence space, using BLOSUM62 as a similarity metric and Laplacian eigenmaps [64] as the dimensionality reduction technique (Fig 3A, first panel). This analysis reveals three distinct sequence clusters: one occupied by the RfaHα pMPNN-SD design results (light blue), one occupied by the RfaHβ pMPNN-SD design results (yellow), and one occupied by the WT RfaH sequence (black), the pMPNN-AD design results (gray), and the genetic algorithm design results (green and purple). This clustering pattern is largely reproduced in the pMPNN-SD and AF2Rank objective space (Fig 3A, second and third panels), although there is a greater spread of sequences in the AF2Rank objective space, which appears to indicate a somewhat greater sensitivity of the AF2Rank composite score to the differential sequence preferences of the two RfaH states (S7 Fig). Nevertheless, the fact that all three embeddings generate similar clustering patterns suggest that the multistate design methods examined so far have all succeeded, to various extent, in recapitulating the WT RfaH sequence profile, which is distinct from hypothetical non-foldswitching single-state RfaH sequence profiles.

**Fig 3.**
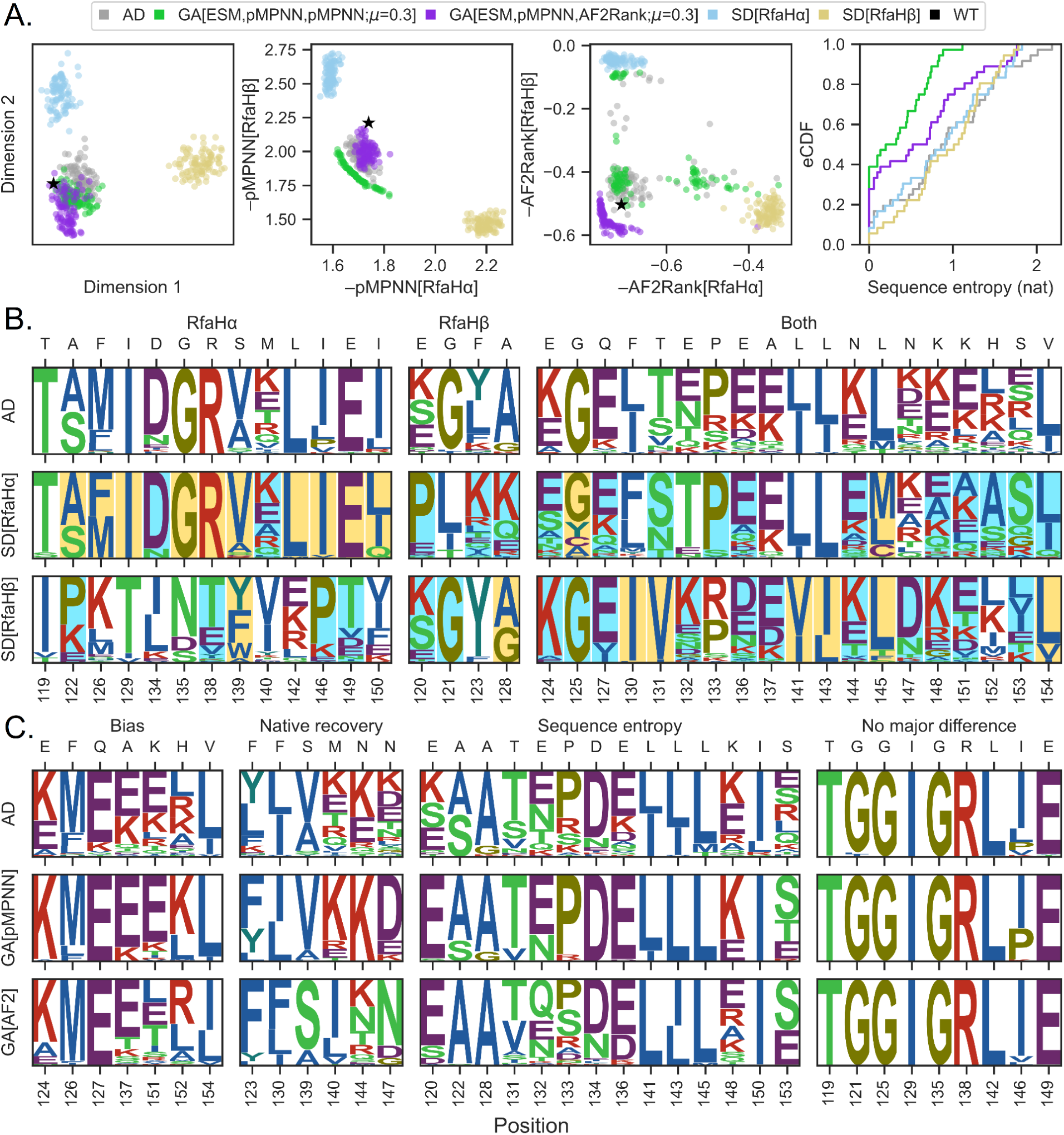
Integration of multiple models into sequence design leads to reduction of sequence bias and variance. A. From left to right: distribution of pMPNN-AD (gray), pMPNN-SD (blue for the RfaHα state and yellow for the RfaHβ state), GA[ESM,pMPNN,pMPNN;*μ*=0.3] (green; later abbreviated as GA[pMPNN]), A[ESM,pMPNN,AF2Rank;*μ*=0.3] (purple; later abbreviated as GA[AF2Rank]), and the WT (black) sequences in a two-dimensional embedding generated with the Laplacian eigenmaps algorithm [64], the pMPNN-SD log likelihood objective function space, and the AF2Rank composite score objective space. The fourth panel on the right shows the empirical cumulative distribution functions (eCDF) for the per-position sequence entropy (base *e*) of these populations of designed sequences. All GA sequences in this figure refer to the final iteration sequence populations. B. Logo plots for sequences generated using pMPNN-AD (top), pMPNN-SD for the RfaHα state (middle), and pMPNN-SD for the RfaHβ state (bottom). The residue positions are organized into three blocks, depending on whether the sequence profiles from the RfaHα (left), RfaHβ (middle), or both states (right) dominate the pMPNN-AD sequence profiles. For the RfaHα and RfaHβ pMPNN-SD design sequence logo plots, the residue positions are shaded according to their relative solvent accessibility in the corresponding state; blue shading indicates a relative solvent accessibility > 50%, while yellow shading indicates < 20%. The WT residue type at each position is indicated on the secondary *x*-axis. C. Logo plots for sequences generated using pMPNN-AD (top), GA[pMPNN] (middle), and GA[AF2Rank] (bottom). The residue positions are organized into four blocks: “bias”, “native recovery”, “sequence entropy”, and “no major difference”; see the main text for more details on this classification.

Next, we investigate the sequence composition underlying the apparent separation between the single-state and multistate design results. This analysis reveals two distinct but equally prevalent patterns in the sequence profiles generated through pMPNN-SD and pMPNN-AD (Fig 3B): conflicting residue positions (first two panels), where the most frequently recommended residue types from one of the two states dominates the multistate sequence profile, and non-conflicting residue positions (third panel), where the multistate sequence profile consists of a union of residue types assigned to both states. In addition, for the 17 positions where the two states have conflicting residue preferences, 13 are dominated by the RfaHα state, and these 13 positions constitute the majority of buried core residues in this state (yellow shading, middle row), while the majority of the RfaHβ core residues (yellow shading, bottom row) are located at non-conflicting positions. Interestingly, of the RfaHα core positions that dominate the multistate sequence preference, all but E149 are located at the N- and C-terminal domain interface. These observations are consistent with the fact that the RfaHα state is highly stabilized when in contact with the N-terminal domain, while the RfaHβ state is the preferred, albeit only marginally stable, state as the two domains dissociate [65]. Overall, this pattern of sequence preference explains the clustering patterns between the single-state and multistate design sequences, and why the multistate design sequences appear to be closer to the RfaHα single-state design sequences in the low-dimensional representation of the RfaH sequence space. Lastly, we note here that, although it is theoretically possible for the multistate sequence profile to be distinct from both single-state designs (e.g., a foldswitching protein may contain a polar residue at a position where its two constituent conformational states may respectively prefer charged and hydrophobic residues), this third scenario is not observed with the current model system.

### Genetic algorithms can reduce bias and variance in native sequence recovery

Having examined closely the difference among the sequences generated using pMPNN-AD and pMPNN-SD, we now turn to the difference among the sequences generated using pMPNN-AD, GA[pMPNN], and GA[AF2Rank]. This leads us to divide the set of designable positions into four groups (Fig 3C): “bias”, where the native residue is recovered at a very low level, if at all, with all three setups; “native recovery”, where GA[pMPNN] and/or GA[AF2Rank] are able to recover the native or native-like residue at significantly higher fraction than pMPNN-AD; “sequence entropy”, where all three methods are able to frequently recover the native residue, but GA[pMPNN] and/or GA[AF2Rank] show lower level of sequence entropy; and “no major difference”, where the three methods have identical or near-identical performance.

Setting aside the “no major difference” category, most of the designable positions differ in terms of sequence entropy in the sequence populations generated using pMPNN-AD, GA[pMPNN], and GA[AF2Rank] (Fig 3C, third column). In particular (Fig 3A, fourth panel), the sequence population generated by GA[pMPNN] (green) tends to have the lowest level of sequence entropy, followed by that generated using GA[AF2Rank] (purple), while the sequences generated using pMPNN-AD (gray) and -SD (light blue and yellow) tend to have the highest level of sequence entropy; this pattern holds for most of the genetic algorithm setups examined in this work (S8 Fig). This reduction in sequence entropy oftentimes concentrates probability on the WT residue types, thus partially explaining why GA[pMPNN] and GA[AF2Rank] have much better native sequence recovery than pMPNN-AD (Fig 2, fifth and sixth columns). Given this observation, we ask whether it is possible to achieve similar sequence entropy reduction by performing a post hoc non-dominated sorting of the pMPNN-AD sequences in the objective spaces; this filtering step results in a significant loss of the sequence population, yet the resulting sequence entropy profiles are no better than that of a random subsampling of the sequence population (S9 Fig), which indicates that the post hoc approximate Pareto front does not confer the same benefit as that generated using true multiobjective optimization.

The “bias” and “native recovery” categories (Fig 3C, first two columns) suggest systematic differences between the RfaH WT sequence and the structure-sequence mapping learned by pMPNN. In particular, at residues Q127, A137, M140, N144, N147, and H152, which are positions that are at least somewhat solvent exposed in one of the two states, pMPNN tends to substitute charged residues for the polar and hydrophobic WT residues, which results in increased surface charges and new surface salt bridge interaction networks (S10 Fig). This is a known bias of pMPNN [20], and explains why the WT sequence is scored somewhat unfavorably by pMPNN-SD (Fig 3A, second panel) compared to the redesigned ones.

At F130, S139, M140, N144, and N147, the WT residues (or, in the case of M140, physicochemically similar residues) are recovered only through GA[AF2Rank]. Among these residues, the effect of charged substitutions such as K and E at M140 and N144 is ambiguous, because such substitutions tend to have lower β-strand propensity, even though an increase in surface charges and salt bridge interactions may improve stability and facilitate strand pairing during folding [66]. On the other hand, F130 and S139 likely play an important role in shaping the free energy landscape of RfaH, in a way that likely cannot be replaced by the mutations suggested by AD and GA[pMPNN]. First, in the RfaHα state, F130 is located between the N- and C-terminal domains and mediate their interface (S11A Fig), and protease accessibility assays have demonstrated that F130V, a substitution to a smaller hydrophobic residue, weakens such interface interaction [67]. Second, S139 is located at α2 in the RfaHα state and β3 in the RfaHβ state. In the RfaHα state, S139 introduces a buried unsatisfied hydrogen bond donor and acceptor into the N- and C-terminal interface (S11C Fig), while in the RfaHβ state its sidechain forms a hydrogen bond interaction with that of N156 near the core of the domain (S11B Fig); as such, a mutation to V or A is expected to further stabilize the RfaHα state at the expense of the RfaHβ state. Therefore, at residue position 139, GA[pMPNN] designs appear to favor the RfaHα state, while the GA[AF2Rank] designs tend to maintain the WT serine residue. However, in the WT, the N-terminal portion of α2 already exhibits high local stability [68] and transient helical content in the foldswitching transition state ensemble [69,65], likely helped by its high leucine content (L141, L142, L143, and L145) [70], while the C-terminal all-β state is only marginally stable, even though it is the energetically preferred state when the C-terminal domain is dissociated from the N-terminal domain [65]. Therefore, by maintaining WT residues, the GA[AF2Rank] designs are more likely to retain the foldswitching phenotype of the WT sequence compared to the GA[pMPNN] designs. These results therefore highlight how integrating multiple models into the sequence design process can produce sequences that may capture key energetic effects better than what each model can produce individually.

### pMPNN hyperparameter tuning leads to bias-variance tradeoff

In the previous section, we demonstrated how the Pareto optimality condition in the objective space can manifest in the sequence space both in terms of bias reduction (i.e., concentration of probability on the WT residue types) and variance/entropy reduction (i.e., concentration of probability on a smaller set of residue types). This result is achieved without any finetuning of pMPNN hyperparameters, such as temperature or state weights; in this section, we analyze whether modifications to these hyperparameters might yield similar improvements without invoking the evolutionary multiobjective optimization framework.

First, we examine whether it is possible to reduce pMPNN-AD sequence entropy by lowering the sampling temperature. To answer this question, we perform additional sequence design with pMPNN-AD at the temperature of 0.1, and compare the results with existing simulations performed at temperature 0.3. As expected, reducing the sampling temperature reduces sequence entropy (S12B Fig, fourth panel), which leads to an improvement in native sequence recovery (S12A Fig). This is manifested in the pMPNN-SD objective function space as a partial overlap between the higher temperature Pareto front and the lower temperature pMPNN-AD population (S12B Fig, second panel). However, in the AF2Rank objective space, lowering the temperature moves the pMPNN-AD population further away from the GA[AF2Rank] Pareto front (S12B Fig, third panel). Consistent with this observation, at this lower temperature, pMPNN-AD no longer recovers the native sequence at positions Q127, S139, and N144, and recovers the native sequence with a much lower probability at positions F123 and F126 (S12C Fig). As such, while lowering the temperature reduces variance, it also increases the bias in the redesigned pMPNN-AD sequences.

Next, we examine the effect of state weights in native sequence recovery. In pMPNN-AD, the model takes a weighted average of the state-dependent logits during autoregressive sequence decoding, with the default weight being 1.0 for each state (before normalizing the sum of all state weights to 1.0). As we saw with Fig 3B, the pMPNN-AD sequence profiles are the results of balancing the oftentimes conflicting sequence preferences of each constituent state.

Therefore, one plausible explanation for why pMPNN-AD and GA[pMPNN] sometimes fail to recover the native sequence is that there is a mismatch between the state weights and the actual local stability difference between the two states of WT RfaH. To assess this hypothesis and whether it can be corrected by rebalancing the state weights, we extract the raw logits (conditioned on the rest of the WT sequence) for positions in the “native recovery” group in Fig 3C, and use the logits to compute how the pMPNN-AD sequence profiles at these positions change as a function of the state weights (S13 Fig). We find that for F130, native recovery could have been significantly improved by a stronger weight for the RfaHα state, although this does come with increased sequence entropy. For S139, M140, N144, and N147, however, varying the state weights would not have meaningfully improved native sequence recovery, because the native residue is never among the preferred residue types under any state weight choices; the fact that GA[AF2Rank] succeeds in recovering the native sequence at many of these positions highlights the ability of the genetic algorithm to pick out and amplify small signals, when equipped with the appropriate objective functions. Taken together, it appears that adjusting state weights is not a useful strategy for correcting biases in native sequence recovery; even in cases such as F130 where the state weight appears to have been misspecified, it is unclear how, a priori, one should set the per-position state weights, especially considering the fact that these weights do not have a straightforward biophysical interpretation.

In conclusion, if sequence design is viewed as a statistical inference problem, the effect of tuning the pMPNN state weights and sampling temperature is subject to the bias-variance tradeoff, where hyperparameter choices that reduce bias come at the cost of increased variance, and vice versa. This is to be contrasted with the results from the previous section, where we showed that GA[pMPNN] and GA[AF2Rank] can reduce bias and/or variance in the population of redesigned sequences, through the integration and Pareto optimization of multiple sources of information on the RfaH sequence-structure relation.

### Genetic algorithms improve sequence similarity to RfaH-like foldswitching proteins

Even though native sequence recovery is a standard metric for assessing the performance of sequence design methods, the metric may have difficulty capturing the sequence patterns underlying the foldswitching capacity, or lack thereof, of the redesigned RfaH sequences. This assessment is based on the estimation that the average pairwise sequence identity of foldswitching sequences related to the NusG superfamily, to which RfaH belongs, is only around 20% [71]. Given this observation, we seek an additional metric to assess how well the RfaH sequences redesigned by the multistate methods might retain the RfaH foldswitching phenotype. To do so, we compare the redesigned sequences to a computationally annotated and experimentally validated database of NusG-like sequences [71], which are predicted to be either foldswitchers or non-foldswitchers (see Methods and Fig 4). We find that sequences designed by GA[pMPNN] and GA[AF2Rank] tend to be more similar to the foldswitcher sequences than those designed using pMPNN-AD, while the similarity to the non-foldswitchers is largely unaffected. However, the extent to which the genetic algorithm outperforms pMPNN-AD depends on the mutation rate, and, again, the mutation rate 0.3 appears to produce the best sequences. These results are consistent with our analysis of the GA[pMPNN] and GA[AF2Rank] sequences so far, and confirm that these methods have the potential to outperform pMPNN-AD for multistate protein sequence design.

**Fig 4.**
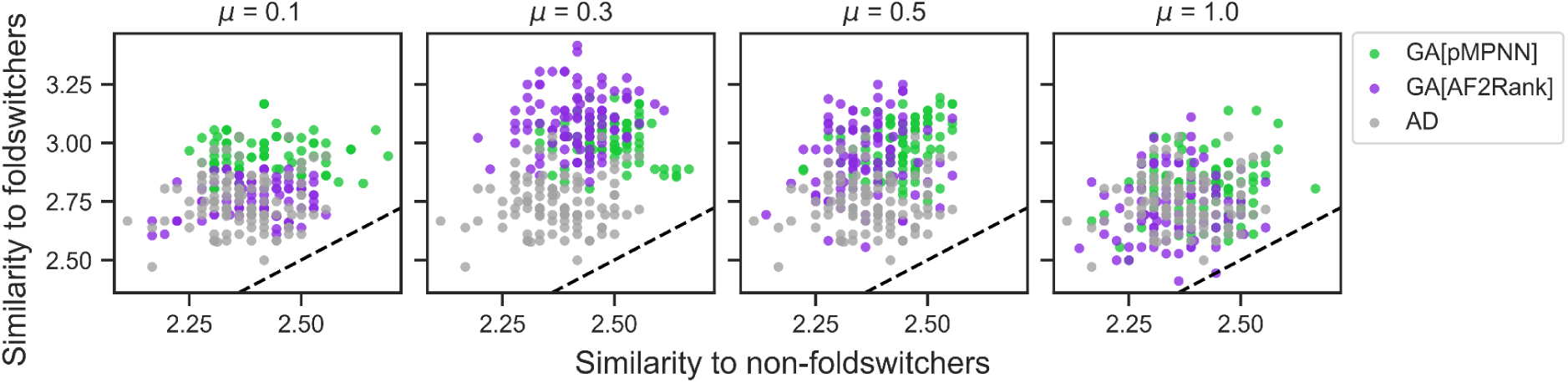
Genetic algorithm designs exhibit greater sequence similarity to NusG-like foldswitching proteins than non-foldswitching proteins. The four panels show the distribution of sequence similarity measures for the pMPNN-AD (gray), GA[pMPNN] (green), and GA[AF2Rank] (purple) sequences to NusG-like foldswitching and non-foldswitching sequences. Each panel corresponds to a different mutation rate for the GA simulations, indicated in the panel title. The dashed black lines represent the *y* = *x* lines. See Methods for more details on the similarity measure and the NusG-like sequence database.

## Discussion

In this work, we examined the potential of evolutionary multiobjective optimization as an integrative framework for protein sequence design. This framework was chosen because of its ability to explicitly approximate the Pareto front in a user-specified objective space, and the flexibility it affords to construct informative mutation operators to guide sampling in the sequence space. Using the multistate design problem of the foldswitching protein RfaH as an in-depth case study, we showed that this approach led to design candidates with reduced bias and variance in native sequence recovery, without the need for post hoc filtering or pMPNN hyperparameter tuning.

Moving forward, as more models become available that capture additional aspects of protein sequence-structure-function relationships, we anticipate such an evolutionary multiobjective optimization framework to be broadly relevant for and readily adaptable to even more complex design tasks. For example, the framework may be adapted for the design of biologics that simultaneously optimizes stability, affinity, specificity, immunogenicity, and pharmacokinetics, or for the construction of entire signaling or metabolic pathways, where multiple biomolecules are simultaneously designed to optimize functional properties. For such design tasks, however, genetic algorithms designed to handle higher-dimensional objective spaces, such as NSGA-III [72], may be more appropriate.

Other studies have employed Monte Carlo sampling to invert models such as AlphaFold2 for design [20,21,43,51,59,60,63,73]. However, such an approach can generate “adversarial” sequences with poor biophysical properties that nevertheless optimize the objective functions [21]. This observation suggests that, despite the innovation in model architecture and availability of curated sequence and structure datasets, it remains challenging to perfectly align the local maxima of a learned sequence-structure mapping with the true protein fitness landscape. Here we argue that the evolutionary multiobjective optimization framework demonstrated in this work is less likely to produce such artifacts, for two reasons. First, when multiple models feed into the generative process, as is the case in this work, an adversarial sequence can be propagated only if it is overfitted to by all models. This is less likely if the models rely on different architectures, loss functions, and training data, and as such their errors are unlikely to be consistently aligned. Second, in a multiobjective optimization setting, the Pareto front often does not contain solutions that simultaneously optimize all objective functions; such solutions are commonly known as either the ideal point or the utopia point, and its absence is indicative of underlying tradeoff conditions in the design space. Taken together, the use of informative mutation operators, in conjunction with the Pareto optimization of the objective functions, may minimize adversarial sequences in a way analogous to the voting ensemble technique [74].

Recently, denoising diffusion probabilistic models [35–44] have emerged as a powerful class of generative models, especially for the purpose of de novo protein backbone generation. Such diffusion models share some similarities to the evolutionary multiobjective optimization framework examined in this work. Both classes of methods transform corrupted samples into biophysically realistic ones through a sequence of intermediate distributions; in a diffusion model, this transformation is achieved through a learned, time-dependent transition kernel, while in this work, this transformation is achieved through a time-independent mutation operator, guided by the Pareto optimality condition. In addition, both classes of methods share an interesting connection with autoregressive models. Some discrete diffusion models can be viewed as a generalization of autoregressive models, when the autoregressive sequence decoding steps are viewed as the reverse of a sequential corruption forward process; however, without the restriction of sequential refinement, such diffusion models have the advantage of a parallel, iterative sampling process [75]. As we demonstrated in the analysis of the GA[RND,pMPNN,pMPNN] setup, by embedding pMPNN into the mutation operator, we effectively extend the autoregressive sequence decoding model of pMPNN into that of a parallel, iterative sampling process as well, although “parallel” in the current context is more appropriately interpreted as a population-level parallelization.

A key property of diffusion models is that they can be used to integrate additional design objectives, whereby the gradient of any differentiable objective function or classifier can be added to the score function to bias the reverse diffusion process. The framework demonstrated in this work imposes no differentiability conditions. More importantly, however, in a diffusion model, these additional objective functions are collapsed into a scalar value, typically by taking a linear combination. This technique, called scalarization, is commonly used to reduce a multiobjective optimization problem into a single-objective one, but gradient-based optimization of such a linear combination can lead to incomplete recovery of the Pareto front, especially when the Pareto front is not convex [46]. In ensemble/population-based Monte Carlo methods such as NSGA-II, scalarization is unnecessary, because the Pareto front is naturally encoded in the population structure of the candidate solutions, but no equivalent structures exist for single-variable Monte Carlo methods such as diffusion models. Therefore, an interesting line of research is to investigate whether diffusion models can be extended into ensemble/population-based Monte Carlo methods and be used to perform controlled generation explicitly conditioned on the Pareto front in an objective space. Ultimately, however, a more general approach is to construct a generative model that can be directly and efficiently conditioned on the free energy landscape defined in a reaction coordinate space; indeed, some recent progress has been made towards solving the inverse of this problem [39,76].

## Methods

### Structure preparation

The starting RfaH structures are taken from the Protein Data Bank. The RfaHα structure is taken from 5OND [77], after deleting the DNA and water molecules and the symmetric copy. The RfaHβ structure is taken from the first model of 2LCL [56], after deleting residues 97–107. The RfaHα structure contains a missing loop at residues 98–117 that connects the N- and C-terminal domains, which is left unmodeled.

The starting structures are relaxed using a custom script with PyRosetta [78]. Specifically, a gentle CA position restraint is applied to the starting structures, which are then relaxed using FastRelax [79,80] in dual space with the `MonomerDesign2019` script and the `beta_nov16_cart` scoring function [81]. For each starting structure, 1000 relaxed structures are generated, and the structure with the lowest total score is chosen for subsequent simulations. Note that the relaxed structures of the two RfaH states are used as inputs to all pMPNN and AlphaFold2 function calls; the input structures are not updated based on any further designed sequences or predicted structures.

Structural visualization is performed with PyMol (version 2.5.4). Relative solvent accessibility calculation on the relaxed structures is performed using the GETAREA webserver [82].

### ProteinMPNN

The code for ProteinMPNN (pMPNN) [20] (version 1.0.1) was accessed via GitHub from https://github.com/dauparas/ProteinMPNN. In this work, we perform sequence design with pMPNN-AD and pMPNN-SD using the vanilla model weights and the default weight of 1.0 for each state; the designable positions are the C-terminal residues 119 to 154. When pMPNN-SD is used as an objective function, the negative log likelihood score of a state is computed from all residue positions and averaged over 5 repeats (`--score_only`, `--num_seq_per_target 5`). Unless otherwise specified, all pMPNN runs are performed at a temperature of 0.3, which is chosen to balance native sequence recovery and sequence diversity; note that the pMPNN-SD log likelihood scores are temperature-independent.

To perform multistate design with pMPNN-AD, we input a combined PDB file containing the structures for all the states of the system, such that the pairwise centroid distance *r_ij_* between the structures of state *i* and *j* satisfies

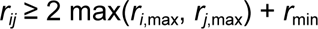

Where *r_i_*, _max_ is the maximum distance between the centroid of structure *i* and the set of all CA atoms in the structure, and *r*_min_ is a minimum distance set to 24 Å. This distance is chosen such that the backbone atoms from the structures of different states will have a shortest distance longer than the range of distances that can be encoded as edge features using the radial basis functions. The corresponding designable positions in the two states of RfaH are then tied together during sequence decoding.

When the sequence encoded in the input structure does not match the sequence of a design candidate, as is almost always the case during genetic algorithm simulations, the sequence in the JSON file generated from parsing the input PDB file is updated before feeding to pMPNN.

### Genetic algorithm

In this work, we employ NSGA-II [55] as the evolutionary multiobjective optimization framework, through its implementation in the `pymoo` package [83] (version 0.6.0.1). Briefly, NSGA-II is an iterative algorithm, and, at each iteration, the population is doubled by the application of binary tournament selection, a crossover operator, and a mutation operator. Then, the new, combined population is partitioned by repeated non-dominated sorting into successive Pareto fronts.

Lastly, the next generation is populated through elitist selection, whereby individuals from successive Pareto fronts are added to the next generation until the population size limit is reached, with a crowding distance sorting mechanism (in the objective space) as a tiebreaker if not all individuals on the last admitted front can be added. A detailed description and analysis of NSGA-II can be found in [55], and a broader survey of evolutionary multiobjective optimization algorithms can be found in [46].

With NSGA-II, we employ the standard binary tournament selection, *n*-point crossover operator, and a custom-constructed mutation operator to generate new candidates, with the crossover operator and the mutation operator applied with a probability of 0.9 and 1.0, respectively. For the *n*-point crossover operator, *n* = 2 unless otherwise specified. The custom mutation operator provides two methods for picking which subset of residue positions to redesign:

1. “Random” (or “RND”): each designable position is considered for redesign with a probability controlled by the mutation rate (*μ*); if no position is chosen after this procedure, a single designable position is picked randomly.
2. “ESM”: a random polypeptide chain containing designable positions is chosen first, and then the residue at each position in the chain is scored by ESM-1v [27] in a single pass with no masking. The designable positions are ranked from the lowest to the highest scores; the first *k* positions with the lowest scores are selected for redesign, where *k* is either 1 or a random number drawn from a binomial distribution Bin(*n*, *μ*), whichever is larger; here, *n* is the total number of designable positions.

Once a subset of design positions is selected, they are mutated through either uniform random sampling from the set of standard amino acids, or through pMPNN-AD. Lastly, the mutated sequences are scored using the pMPNN-SD negative log likelihood scores, the ESM-1v log likelihood scores, and/or the AF2Rank composite scores for each state; note that the sign for each objective function is chosen such that the simulation can be framed as a minimization problem. To accelerate the simulation, options are implemented to parallelize the evaluation of the mutation operator and the objective functions through either MPI (using `mpi4py` [84]) or a job scheduler.

In this work, we generate 100 sequences for each pMPNN-AD or pMPNN-SD benchmark condition, and perform simulations over 50 iterations with a population size of 100 for each genetic algorithm benchmark setup. Note that for all genetic algorithm benchmark simulations, the sequence at the designable positions are fully randomized in the initial population.

To investigate the reproducibility of sequence populations generated by the genetic algorithm, we repeat simulations for the GA[ESM,pMPNN,AF2Rank] setup with four different mutation rates using a different starting random seed. Comparison of the two sets of simulation results suggest a high degree of reproducibility (S14 Fig).

### AF2Rank

The AF2Rank composite score is used in this work as a folding propensity metric. The method is described in [58] and the code was accessed from https://github.com/jproney/AF2Rank.

Briefly, the sequence(s) for each state is provided to AlphaFold2 [14] via the `ColabDesign` package (version 1.1.1; code was accessed from https://github.com/sokrypton/ColabDesign); here, a state can include binding partners, even if they are not redesigned, and residues missing from the structure of the state are trimmed from the input. AlphaFold2 is called with 1 recycle using the `model_1_ptm` parameter set (an option is also available to call `model_1_multimer_v3` [57] if more than one polypeptide chain is present). The structure for the state is provided as a template, after replacing the template sequence with gap tokens, deleting the sidechain atoms, and imputing the CB positions for glycine residues. No multiple sequence alignment is provided. A final composite score is defined as the product of average AlphaFold2 pLDDT (scaled from 0 to 1), AlphaFold2 pTM, and a TM score between the template and AlphaFold2 predicted structure. The AF2Rank composite score is therefore bounded between [0, 1], with a higher value being indicative of greater folding propensity.

Recently, several works have suggested that, by manipulating the input multiple sequence alignment, AlphaFold2 is sometimes able to predict alternative protein conformations [85–87]. We do not consider these techniques in this work, because their need for evolutionary information may restrict their applicability in a protein design pipeline where no such information is available.

### ESM-1v

The ESM-1v pre-trained protein language model [27] (`esm1v_t33_650M_UR90S_1`) is used in this work as a mutational effect predictor. Through a single forward pass with no masking, the model is used either to rank residue positions, or the per-position score is averaged to give the (pseudo)likelihood score for a sequence. It has been shown that this scoring method gives results that are highly correlated with the masked marginal probabilities generated by sequential masking of single positions, while only requiring a single pass of the model [88]. All calculations involving ESM-1v are done using the `pgen` package (version 0.2.3) [88,89], which is available at https://github.com/seanrjohnson/protein_gibbs_sampler.

### Hypervolume

Hypervolume is a standard metric used to assess the area/volume of the objective space, with respect to a reference point, covered by the approximate Pareto front generated using a genetic algorithm [62]. Therefore, the farther away the solutions are from the reference point, and the wider they are spread out over the approximate Pareto front, the higher the hypervolume. In this work, for RfaH, we use (0, 0) as the reference point to measure hypervolume in the (negative) AF2Rank composite score subspace, and, somewhat arbitrarily, (4, 4) as the reference point to measure hypervolume in the pMPNN-SD (negative) log likelihood score subspace. All hypervolume calculations are performed using `pymoo`.

We note here that the apparent bias of AF2Rank against the RfaHα state in the pMPNN-SD designs (Fig 2, last row) is an artifact, due to the fact that the AF2Rank score is calculated for the full protein structure of each state, but the N-terminal domain is absent in the structure of the RfaHβ state. As a result, the AF2Rank composite score of the RfaHα state has a higher lower bound than that of the RfaHβ state, and a single-state design simulation that optimizes the RfaHα state (at the expense of the RfaHβ state) will therefore have a lower hypervolume.

### Sequence analysis

Native sequence recovery is defined as the fraction of designable positions where the redesigned sequence is identical to the WT sequence. We report the population average native sequence recovery as a performance metric.

(Normalized) sequence similarity is calculated with the BLOSUM62 substitution matrix [90] over the designable positions using Biopython [91]; the per-position scores are averaged to produce the sequence similarity metric. The pairwise sequence similarity matrix is used as a precomputed affinity matrix for Laplacian Eigenmaps dimensionality reduction [64] using `scikit-learn` (version 1.1.3) with the default settings [92].

Visualizations of multiple sequence alignments are generated using Logomaker [93]; the color scheme follows that of [94].

### Comparison to NusG-like sequences

We compare the redesigned RfaH sequences to a database of NusG-like sequences that have been annotated by [71] as either foldswitching or non-foldswitching proteins.

We first filter out sequences in the NusG database where the foldswitching prediction confidence is low, and then randomly sample 1,000 each of foldswitching and non-foldswitching sequences, out of the 15,195 sequences in total in the database, for further analysis. For each selected sequence, a new C-terminal domain sequence is defined, because [71] uses a different definition of C-terminal domain than the RfaH designable positions defined in this work. To do so, the full sequences are downloaded (in December 2023) from UniProtKB [95] using their UniProt IDs, and aligned with the *E. coli* RfaH sequence using Clustal Omega [96,97]; for each aligned sequence, residue positions that do not correspond to *E. coli* RfaH residues 119 to 154 are discarded, and the aligned sequence itself is discarded if its new C-terminal domain sequence contains gap(s). In the end, this results in a reduced set of 645 high-confidence, fully alignable, foldswitching NusG-like sequences, and 956 non-foldswitching ones.

The distance between a redesigned RfaH sequence and the reduced set of (non-)foldswitching sequences is defined as the 95% percentile of normalized BLOSUM62 sequence similarity between the redesigned RfaH sequence and each sequence in the reduced set.

### Data and code availability

All code for methods described in this work and for generating and analyzing the benchmark data can be accessed at https://github.com/luhong88/int_seq_des.

## Acknowledgements

This work was supported by a grant from the National Institutes of Health (R35 GM145236 to T.K.). T.K. is a Chan Zuckerberg Investigator. Computations were performed on resources provided by the Wynton HPC at the University of California, San Francisco.

## Supporting information captions

S1 Fig. Genetic algorithm benchmark simulations at additional mutation rates.

S2 Fig. Benchmark of the n-point crossover operator.

S3 Fig. Random resetting operator results in designs with high sequence entropy.

S4 Fig. GA[RND,pMPNN,pMPNN] produces Pareto-optimal sequences that separate from the pMPNN-AD sequences in the objective space.

S5 Fig. Using ESM-1v as an objective function biases sampling towards RfaHβ-like sequences with lower native sequence recovery.

S6 Fig. Pairwise correlation analysis of performance metrics.

S7 Fig. AF2Rank is sensitive to the differential sequence preferences of the two RfaH states.

S8 Fig. Sequence entropy distributions.

S9 Fig. Post hoc non-dominated sorting does not lead to significant sequence entropy reduction.

S10 Fig. GA[pMPNN] introduces new surface charged residues and salt bridges.

S11 Fig. Mutations at F130 and S139.

S12 Fig. Comparison of GA[pMPNN] and GA[AF2Rank] to low temperature pMPNN-AD sequences.

S13 Fig. Effect of state weights on pMPNN-AD sequence decoding.

S14 Fig. Effect of starting random seed on simulation results.

