## Supplementary Information for "An integrative approach to protein sequence design through multiobjective optimization"

### Table of contents

- Supplementary Figures 1–14

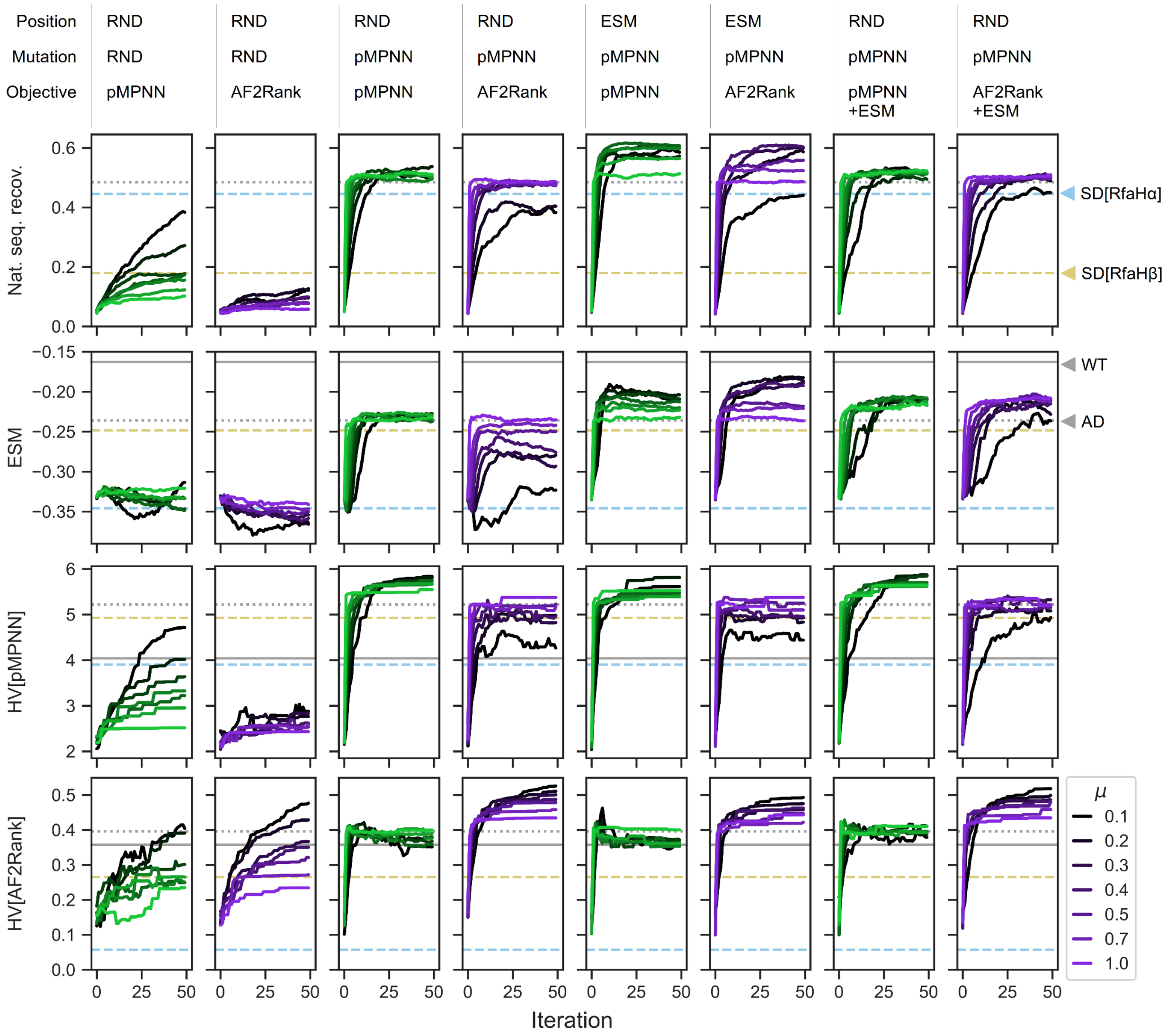


**S1 Fig. Genetic algorithm benchmark simulations at additional mutation rates.** Refer to Figure 2 legend for more details.

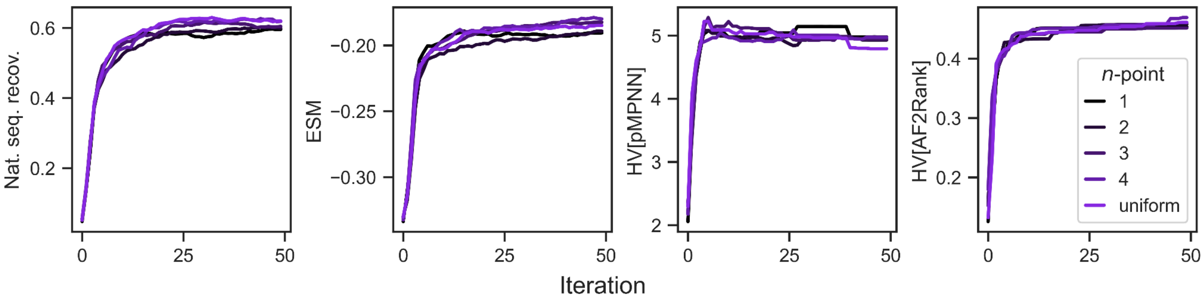


**S2 Fig. Benchmark of the *n*-point crossover operator.** Five sets of simulations are performed with the GA[ESM,pMPNN,AF2Rank;$\mu$=0.3] setup, while varying the number of crossover points. For the uniform crossover operator, the residue at each designable position is chosen from either parental sequences with equal probability; effectively, for $L$ designable positions, the uniform crossover operator is analogous to the ($L$−1)-point crossover operator.

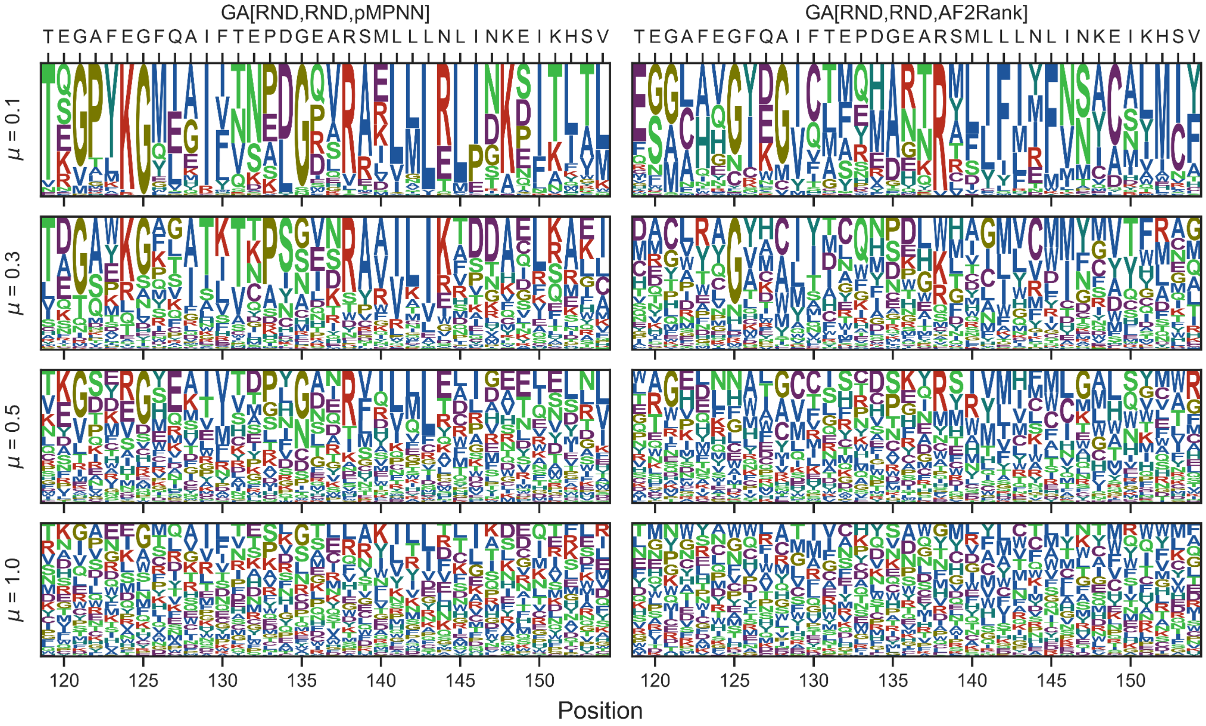


**S3 Fig. Random resetting operator results in designs with high sequence entropy.** Each panel shows the sequence profiles from the last iteration of the GA[RND,RND,pMPNN] (left column) or the GA[RND,RND,AF2Rank] (right column) setup at a different mutation rate (rows). See also SI Figure 8 for the cumulative distribution functions of the per-position sequence entropy.

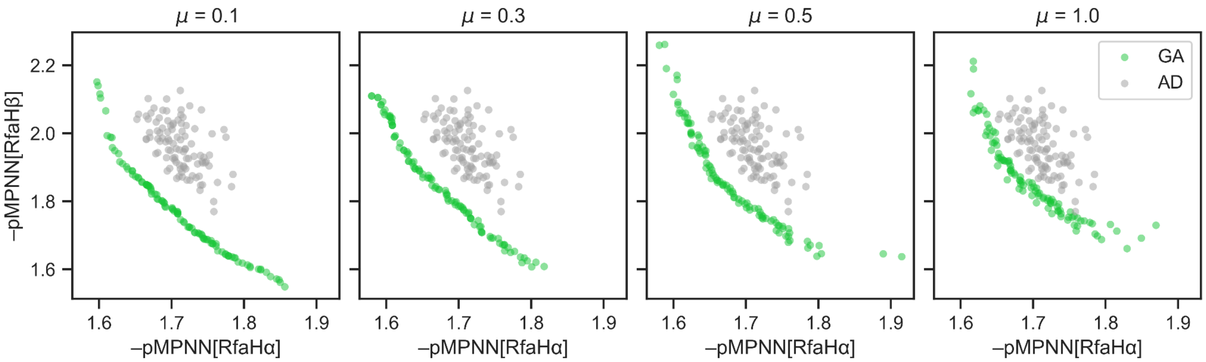


**S4 Fig. GA[RND,pMPNN,pMPNN] produces Pareto-optimal sequences that separate from the pMPNN-AD sequences in the objective space.** Each panel shows the distribution of the last iteration sequences generated using GA[RND,pMPNN,pMPNN] at a different mutation rate (green), compared to the sequences generated from pMPNN-AD (gray), in the pMPNN-SD log likelihood objective function space.

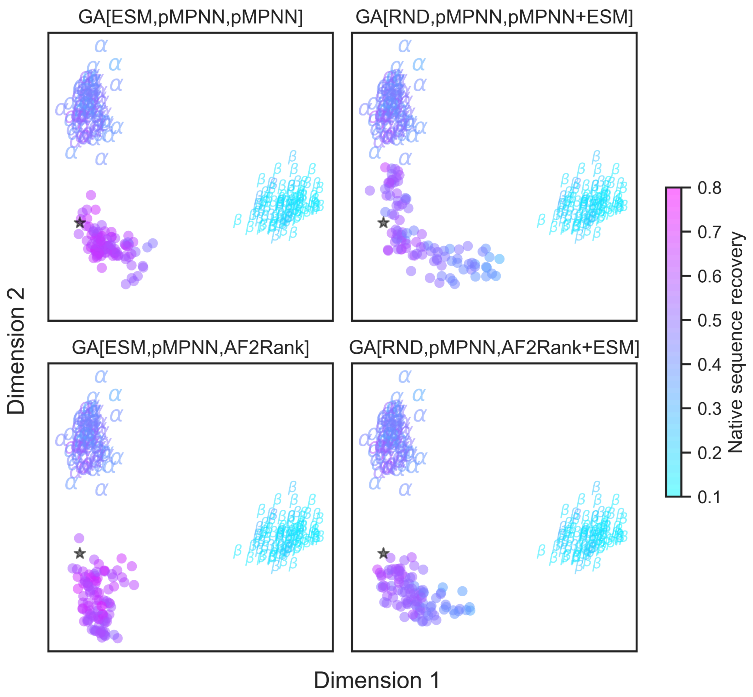


**S5 Fig. Using ESM-1v as an objective function biases sampling towards RfaHβ-like sequences with lower native sequence recovery.** Each panel shows the distribution of the population of the pMPNN-SD sequences and the last iteration population from a GA setup (specified in the title; all with mutation rate 0.3) in a two-dimensional embedding of the sequence space generated using Laplace eigenmaps, as in Figure 2. The sequences are colored by native sequence recovery, except for the WT sequence shown as the black stars; the pMPNN-SD sequences for the RfaHα and RfaHβ states are shown using "*α*" and "*β*" as markers.

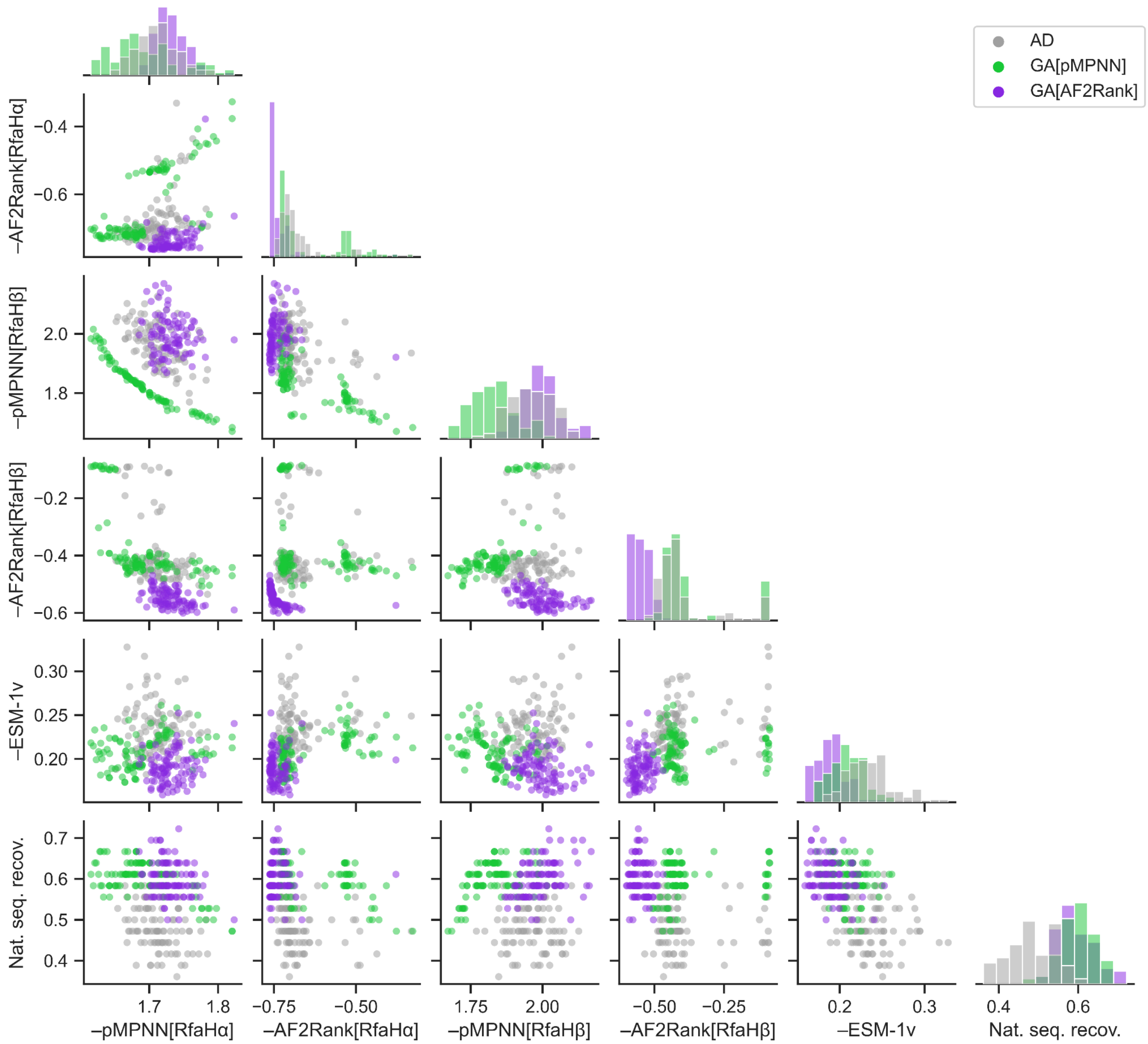


**S6 Fig. Pairwise correlation analysis of performance metrics.** The sequences shown are designed with pMPNN-AD (gray), GA[ESM,pMPNN,pMPNN;$\mu$=0.3] (green; abbreviated as GA[pMPNN]), and GA[ESM,pMPNN,AF2Rank;$\mu$=0.3] (purple; abbreviated as GA[AF2Rank]). All GA sequences in this figure refer to the final iteration sequence populations.

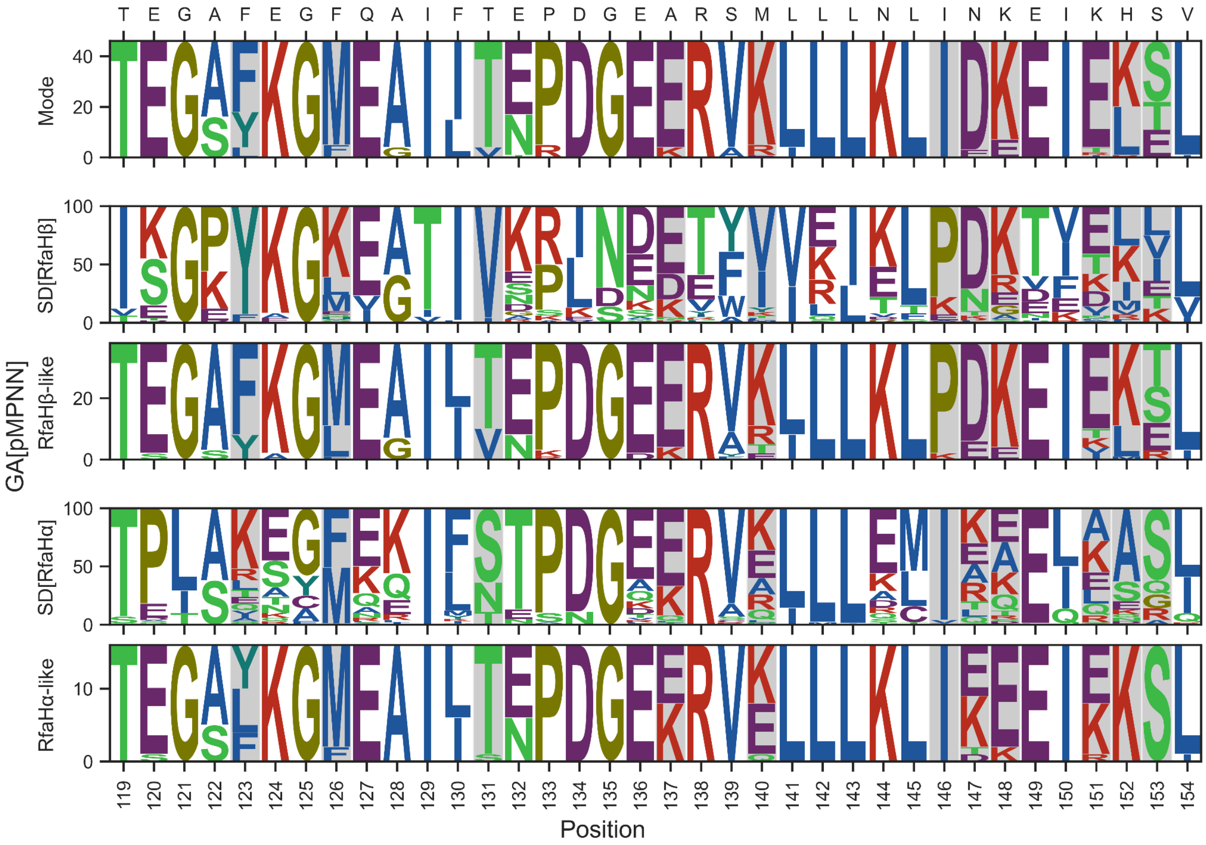


**S7 Fig. AF2Rank is sensitive to the differential sequence preferences of the two RfaH states.** The logo plots for the final iteration sequences designed by GA[ESM,pMPNN,pMPNN;*μ*=0.3] (i.e., GA[pMPNN]), which are partitioned here into three groups: “RfaHα-like”, which has bad AF2Rank[RfaHβ] scores (< 0.6); “RfaHβ-like”, which has bad AF2Rank[RfaHα] scores (< 0.3); and “mode”, which consists of the rest of the sequences. See Figure 3A (third panel) for the distribution of these sequences in the AF2Rank objective space. The logo plots for the RfaHα and RfaHβ single-state design (SD) sequences are also shown for comparison. The positions highlighted in gray shading indicate major differences in the recovered residue types among the three subgroups of GA[pMPNN] sequences.

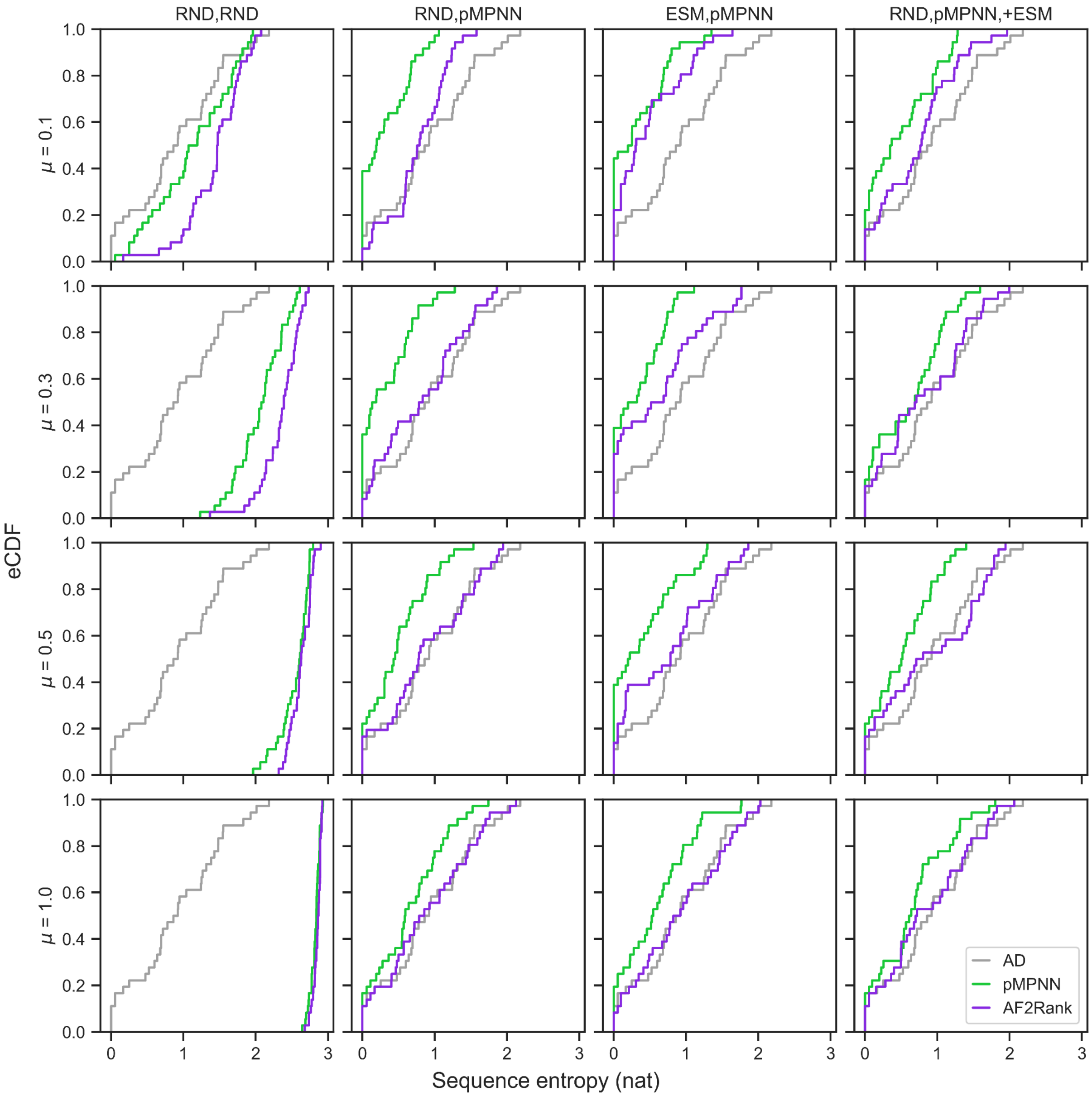


**S8 Fig. Sequence entropy distributions.** Each panel shows the empirical cumulative distribution function (eCDF) of per-position sequence entropy (base *e*) for the last iteration population generated with two genetic algorithm setups. Each row represents a mutation rate ($\mu$), and each column represents a mutation operator configuration ("+ESM" indicates that ESM-1v is used as a third objective function); the curves in green indicate that pMPNN-SD log likelihood is used as the objective functions, and the curves in purple indicate that AF2Rank composite score is used as the objective functions. The gray curves are calculated from the pMPNN-AD reference population. See Figure 2 legend for details on the abbreviations used here.

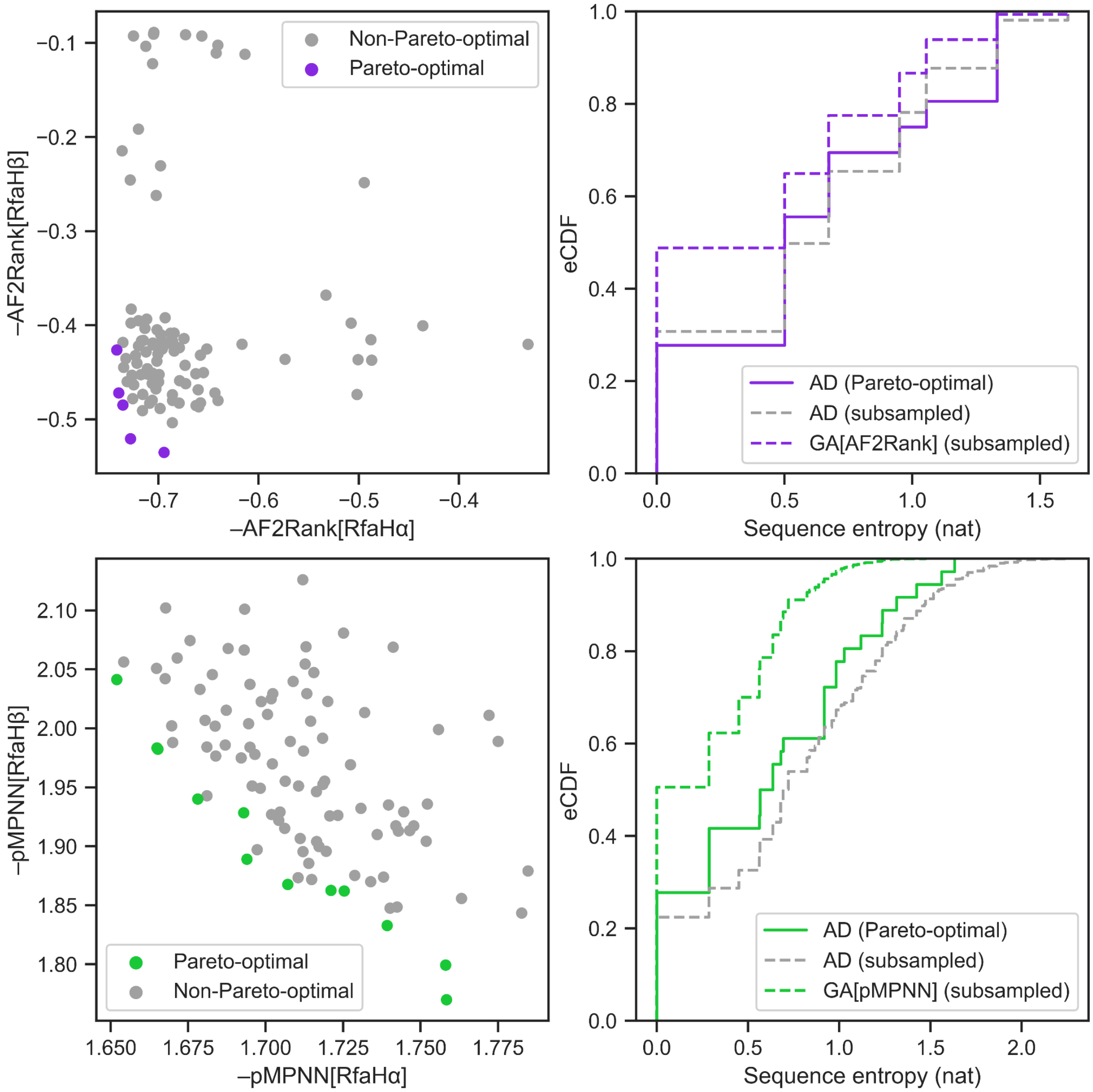


**S9 Fig. Post hoc non-dominated sorting does not lead to significant sequence entropy reduction.** Non-dominated sorting of the 100 pMPNN-AD sequences is performed in the AF2Rank (top row) and pMPNN-SD (bottom row) objective spaces, and the sorting results are shown on the left column. The empirical cumulative distribution functions (eCDF) of the sequence entropies for the sorted, Pareto-optimal solutions are shown on the right column as colored continuous curves. For comparison, sequence entropies are also calculated for a random subsample of the pMPNN-AD sequences of the same size as the Pareto front; this procedure is repeated 100 times and the cumulative distributions of all subsampled sequence entropies is shown as the gray dashed curves. A similar random subsampling method is applied to the GA[AF2Rank] and GA[pMPNN] sequences, and the results are shown as colored dashed curves.

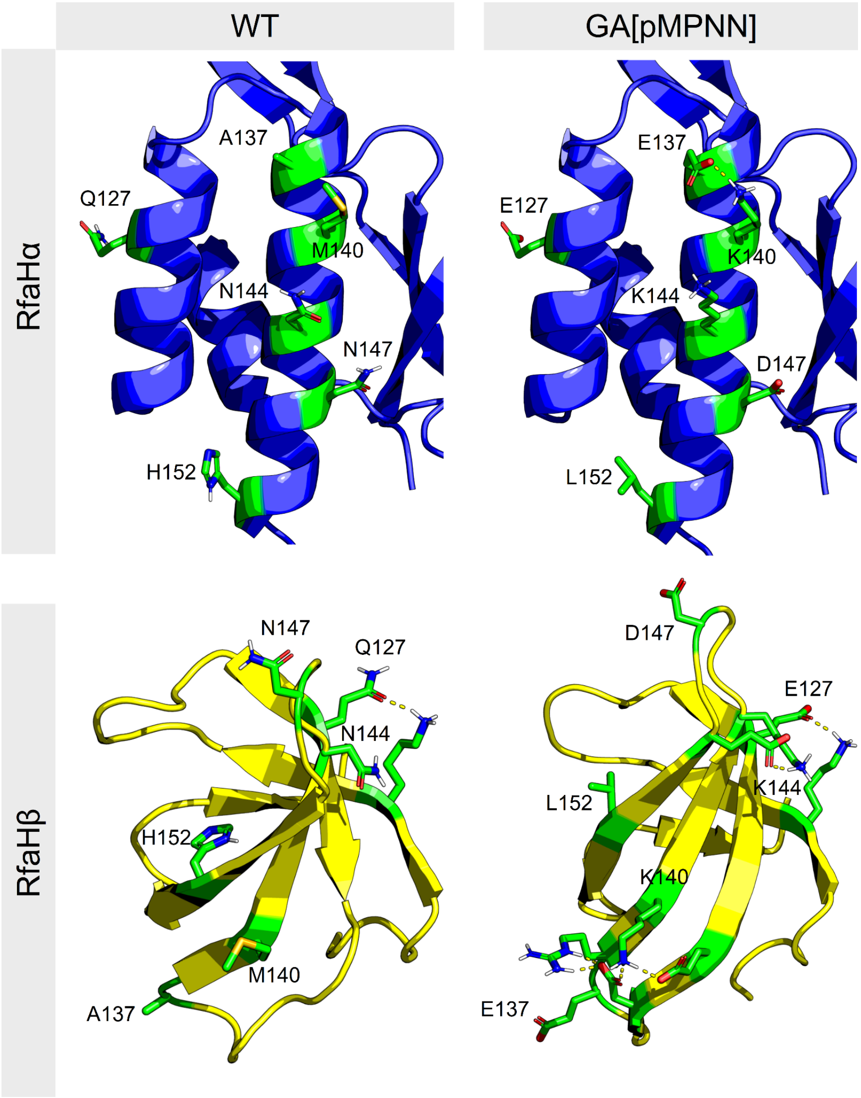


**S10 Fig. GA[pMPNN] introduces new surface charged residues and salt bridges.** The structure models of the WT (left) and a redesigned (right) sequence in the RfaHα (top) and RfaHβ (bottom) states are shown in cartoon representation. The sidechain conformations at the positions 127, 137, 140, 147, 152, and any undesigned positions that form salt bridge interactions with these residues, are highlighted in green stick representation; the salt bridge interactions are represented as dashed yellow lines. The redesigned sequence is chosen as the sequence closest (in terms of percentage sequence identity) to the consensus sequence of the last iteration population of the GA[ESM,pMPNN,pMPNN;$\mu$=0.3] simulation, and the structure models are generated from the relaxed WT structures in the same way as the relaxed WT structures are generated from the experimental structures (see Methods), except that only 10 mutant structures are generated for each state.

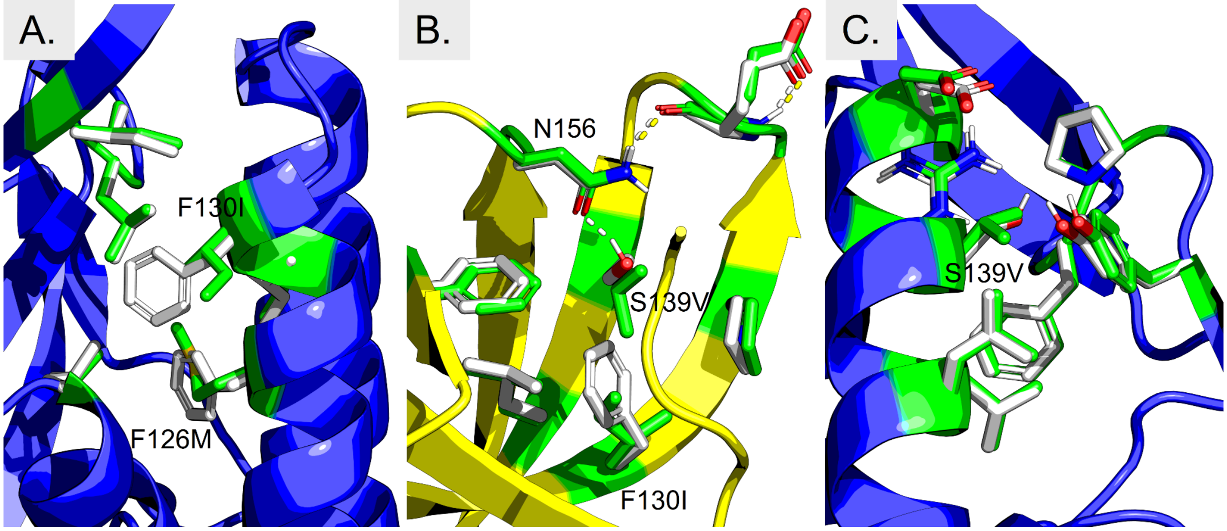


**S11 Fig. Mutations at F130 and S139.** The local structural environment around F130I in the RfaHα state is shown in A, and the local environment around S139V in the RfaHβ and RfaHα states are shown in B and C, respectively. The backbone of the redesigned structure is shown in cartoon representation, while the sidechains of the WT and redesigned residues are shown in white and green stick representation, respectively. The hydrogen bonding interactions are shown with dashed lines, with those in the WT structure in white and those in the redesigned structure in yellow. See SI Figure 10 for more details on how the redesigned sequence is selected and how its structure models are generated.

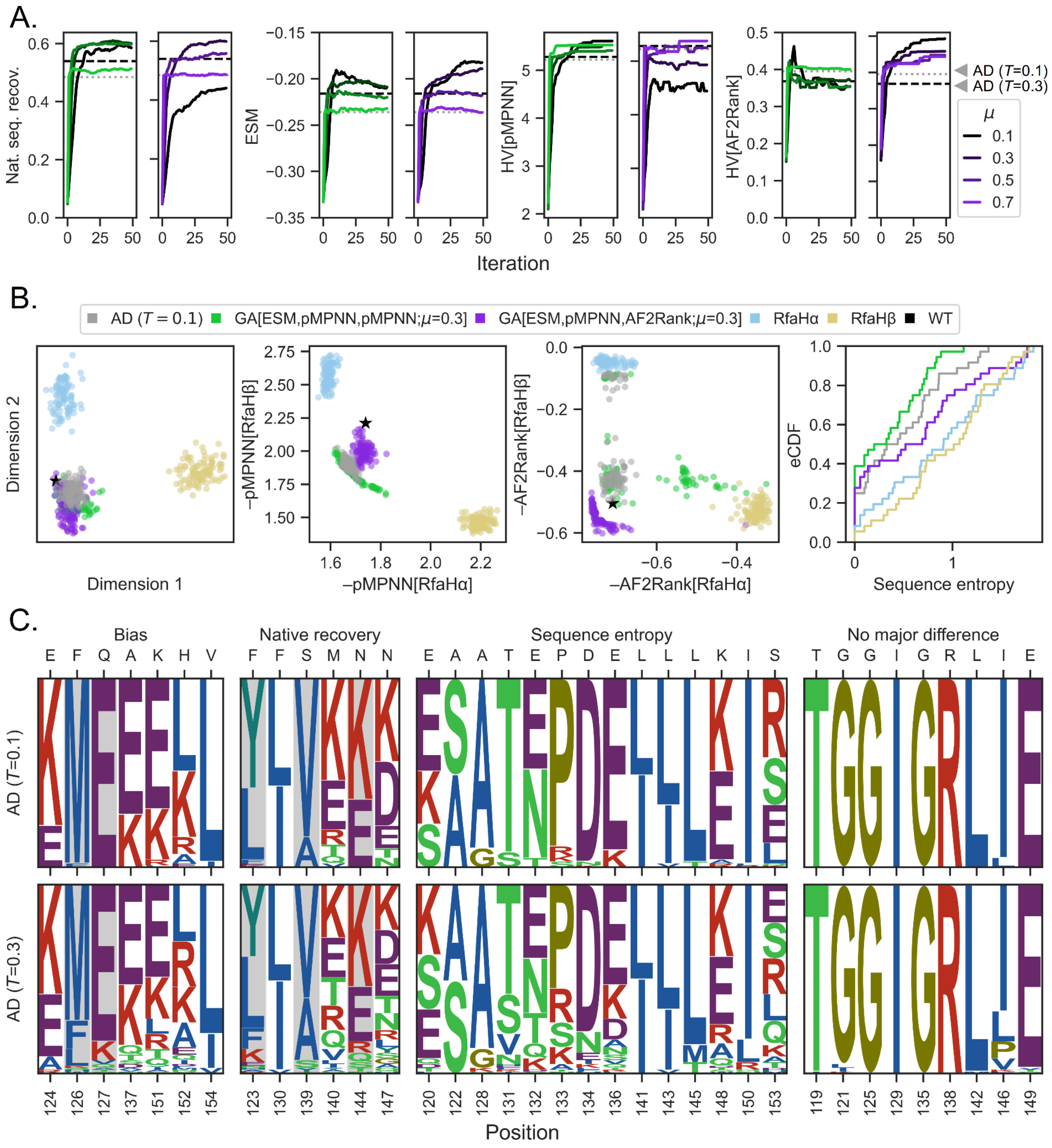


**S12 Fig. Comparison of GA[pMPNN] and GA[AF2Rank] to low temperature pMPNN-AD sequences.**

A. Progression of GA[ESM,pMPNN,pMPNN] and GA[ESM,pMPNN,AF2Rank] simulations, reproduced from Figure 2. The dashed black lines represent the population averages from the pMPNN-AD sequences at $T$ = 0.1, while the dotted gray line represents averages at $T$ = 0.3, as per Figure 2.
B. Distribution of pMPNN-SD, pMPNN-AD, and GA sequences, reproduced from Figure 3A, but with the pMPNN-AD population from the $T$ = 0.1 simulation.
C. Logo plots for sequences generated with pMPNN-AD at $T$ = 0.1 (top) and $T$ = 0.3

(bottom). The residue positions highlighted in gray shading show reduced or no native sequence recovery at *T* = 0.1 compared to *T* = 0.3.

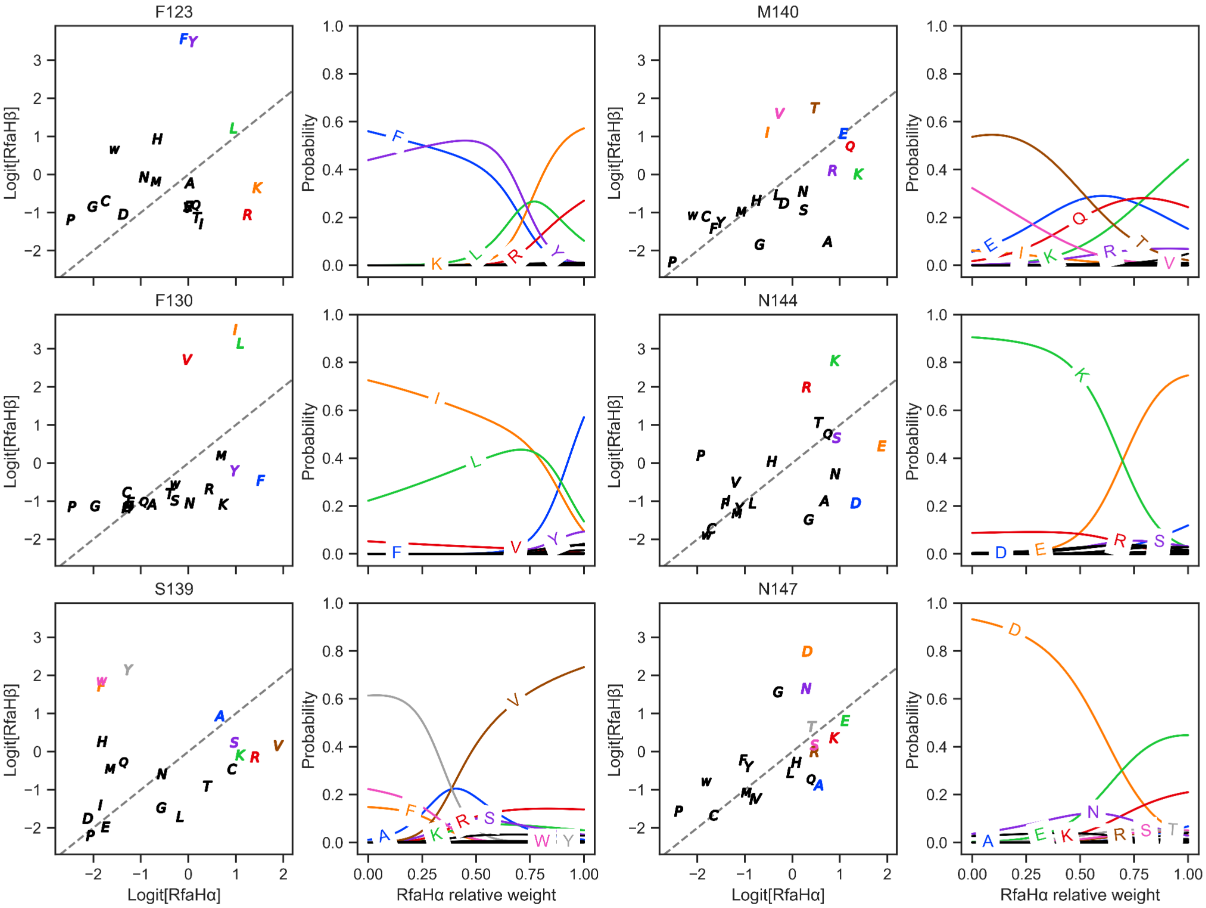


**S13 Fig. Effect of state weights on pMPNN-AD sequence decoding.** For F123, F130, S139, M140, N144, and N147 (i.e., positions identified in Figure 3C in the “native recovery” category), we perform a single-position sequence design simulation with pMPNN-AD to extract the raw logits (not scaled by temperature or normalized by softmax) of all 20 standard residue types for the two RfaH states (first and third columns). The dashed gray lines represent the $y = x$ diagonal lines. Then, we ask how the probability assigned to each residue type (at temperature 0.3) changes as a function of the relative weight for the RfaHα state (second and fourth columns). If a residue type is ever assigned a probability > 0.05 as a function of state weight at a position, then the residue is shown in color on the panels associated with the position; otherwise the residue is shown in black. Note that the relative proportions of decoded residues using pMPNN-AD shown in Figure 3 will not necessarily match those at relative weight 0.5 shown here, because in practice sequence decoding at any given position is usually not performed with the full WT sequence context.

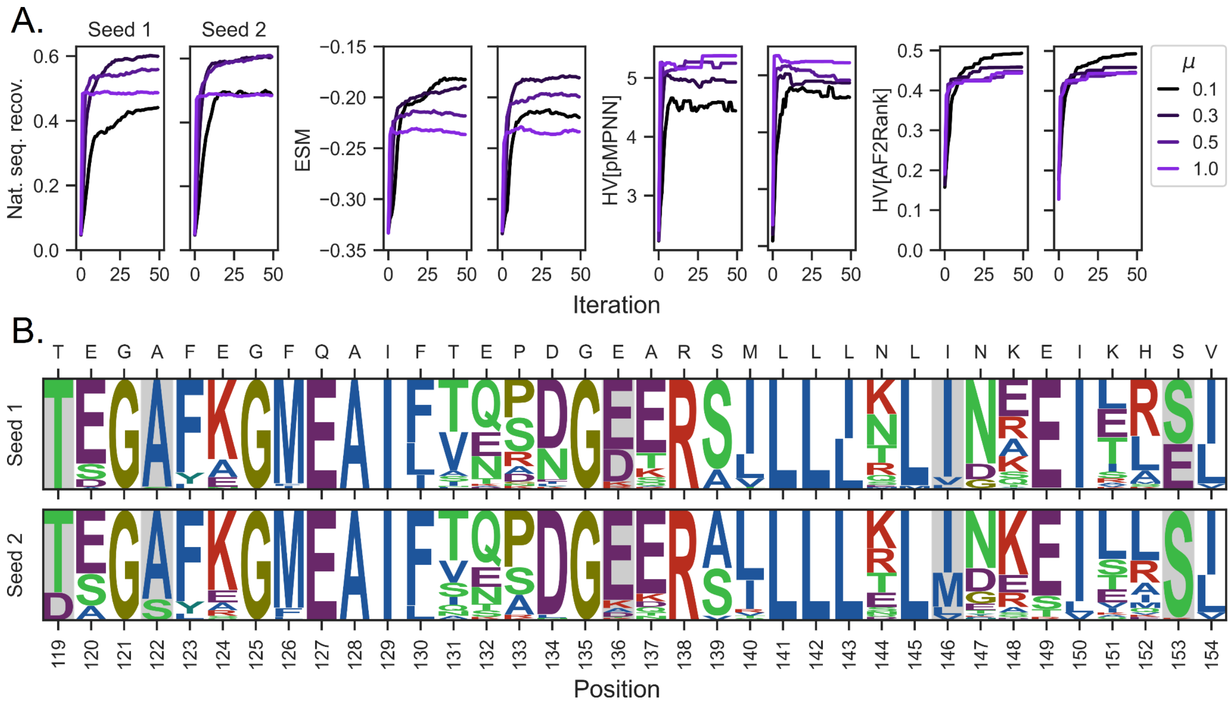


**S14 Fig. Effect of starting random seed on simulation results.**

A. Comparison of convergence metrics for GA[ESM,pMPNN,AF2Rank] simulations at four mutation rates using two different starting random seeds. See Figure 2 for more details on the metrics. The simulation results presented in the rest of this work is based on seed 1.

B. Comparison of the last iteration sequence logos at the mutation rate $\mu$ = 0.3 for the two random seeds. Positions with major differences in the recovered sequence profiles are highlighted in gray shading; in both simulations, the WT residue types are recovered at these positions, but the simulations differ in terms of the alternative residue types recovered at these positions.
